# 3D Reconstruction of Bird Flight Trajectories Using a Single Video Camera

**DOI:** 10.1101/340232

**Authors:** M.V. Srinivasan, H.D. Vo, I. Schiffner

## Abstract

Video cameras are finding increasing use in the study and analysis of bird flight over short ranges. However, reconstruction of flight trajectories in three dimensions typically requires the use of multiple cameras and elaborate calibration procedures. We present an alternative approach that uses a single video camera and a simple calibration procedure for the reconstruction of such trajectories. The technique combines prior knowledge of the wingspan of the bird with a camera calibration procedure that needs to be used only once in the lifetime of the system. The system delivers the exact 3D coordinates of the position of the bird at the time of every full wing extension and uses interpolated height estimates to compute the 3D positions of the bird in the video frames between successive wing extensions. The system is inexpensive, compact and portable, and can be easily deployed in the laboratory as well as the field.

## 1. Introduction

The increasing use of high-speed video cameras is offering new opportunities as well as challenges for tracking three-dimensional motions of humans and animals, and of their body parts (e.g. Shelton et al., 2014; Straw et al., 2011; Fontaine et al., 2009; Dakin et al., 2016); Ros et al., 2017; Troje, 2002; de Margerie et al., 2015; Jackson et al., 2016; Macfarlane et al., 2015; Deetjen et al., 2017).

Stereo-based approaches that use two (or more) cameras are popular, however they require (a) synchronisation of the cameras (b) elaborate calibration procedures (e.g. Hedrick, 2008; Hartley and Zisserman, 2003; Theriault et al., 2014; Jackson et al., 2016) (b) collection of large amounts of data, particularly when using high frame rates; and (c) substantial post-processing that entails frame-by-frame tracking of individual features in all of the video sequences, and establishing the correct correspondences between these features across the video sequences (e.g. Cavagna et al., 2008). This is particularly complicated when tracking highly deformable objects, such as flying birds.

Vicon-based stereo trackers simplify the problem of feature tracking by using special reflective markers or photodiodes attached to the tracked (e.g. Ros et al., 2017; Goller and Altshuler, 2014; Tobalske et al., 2007; Troje, 2002). However, these markers can potentially disturb natural movement and behaviour, especially when used on small animals.

A novel recent approach uses structured light illumination produced by a laser system in combination a high-speed video camera to reconstruct the wing kinematics of a freely flying parrotlet at 3200 frames/second (Deetjen et al., 2017). However, this impressive capability comes at the cost of some complexity and works best if the bird possesses a highly reflective plumage of a single colour (preferably white).

GPS-based tracking methods (e.g. Bouten et al., 2013) are useful for mapping long-range flights of birds, for example, but are not feasible in indoor laboratory settings, where GPS signals are typically unavailable or do not provide sufficiently accurate positioning. Furthermore, they require the animal to carry a GPS receiver, which can affect the flight of a small animal.

A simple technique for reconstructing 3D flight trajectories of insects from a single overhead video camera involves tracking the position of the insect as well as the shadow that it casts on the ground (e.g. Zeil, 1993; Srinivasan et al., 2000). However, this technique requires the presence of the unobscured sun in the sky, or a strong artificial indoor light, which in itself could affect the animal’s behaviour. (The latter problem could be overcome, in principle, by using an infrared source of light and an infrared-sensitive camera).

This paper presents a simple, inexpensive, compact, field-deployable technique for reconstructing the flight trajectories of birds in 3D, using a single video camera. The procedure for calibrating the camera is uncomplicated and is an exercise that needs to be carried out only once in the lifetime of the lens/camera combination, irrespective of where the system is used in subsequent applications.

The system was used in a study of bird flight (Vo et al., 2016) but that paper provided only a cursory description of the technique. This paper provides a comprehensive description of an improved and extended version of the underlying technique and procedure, which will enable its application to other laboratory and field studies of bird flight.

## 2. Methodology

### 2.1 Derivation of basic method

Our method uses a single, downward-looking camera positioned at the ceiling of the experimental arena in which the birds are filmed. The camera must have a field of view that is large enough to cover the entire volume of space within which the bird’s flight trajectories are to be reconstructed.

Essentially, the approach involves combining knowledge of the bird’s wingspan (which provides a scale factor that determines the absolute distance of the bird from the camera) with a calibration of the camera that uses a grid of known geometry drawn on the floor. This calibration provides a means of accounting for all of the imaging distortions that are introduced by the wide-angle optics of the camera lens. In our initial derivation, we assume that the bird does not display a significant amount of roll during its tracked flight. In other words, the two wingtips are in the horizonal plane (or approximately so) when the wings are fully extended. In the Supplementary Information (Section B) we derive and validate a method that accounts for the effects of roll, and also measures the roll angle.

A square grid of known mesh dimensions is laid out on the floor. The 2D locations (X,Y) of each of the intersection points are therefore known. Figure 1 illustrates, schematically, a camera view of the grid on the floor, and of a bird in flight above it, as imaged in a video frame in which the wings are fully extended.

**Figure 1.**
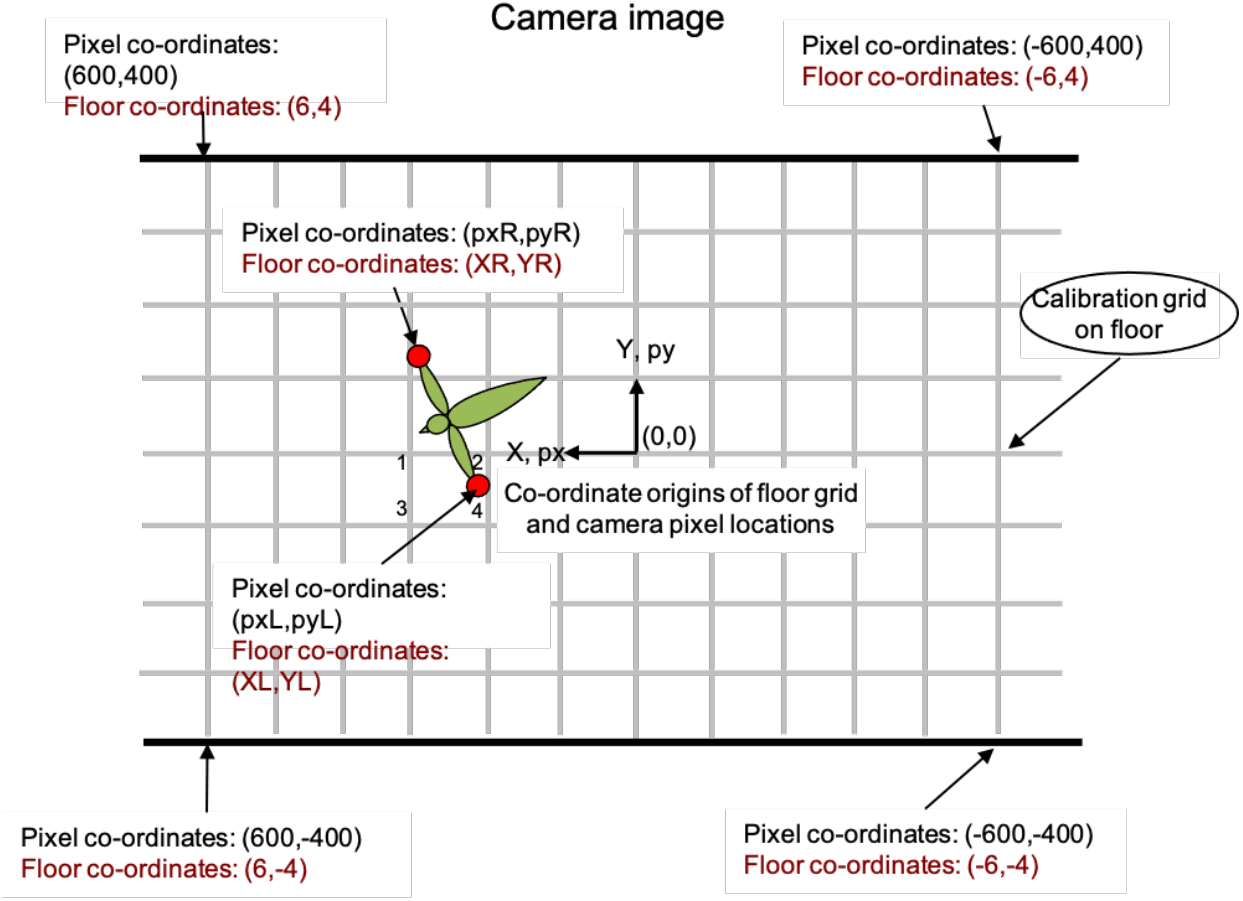
Schematic view of image of the flight chamber from an overhead video camera, showing the calibration grid on the floor, and the instantaneous position of a bird with its wings extended. The origin of the pixel co-ordinates is taken to be the center of the image, i.e. corresponding to the direction of the camera’s optic axis. The origin of the calibration grid is taken to be the point directly beneath the camera, i.e. the position where the optic axis of the camera intersects the floor.

In general, the image of the grid will not be square, but distorted by the non-linear off-axis imaging produced by the wide-angle lens, as shown in the real image of Figure 2.

**Figure 2.**
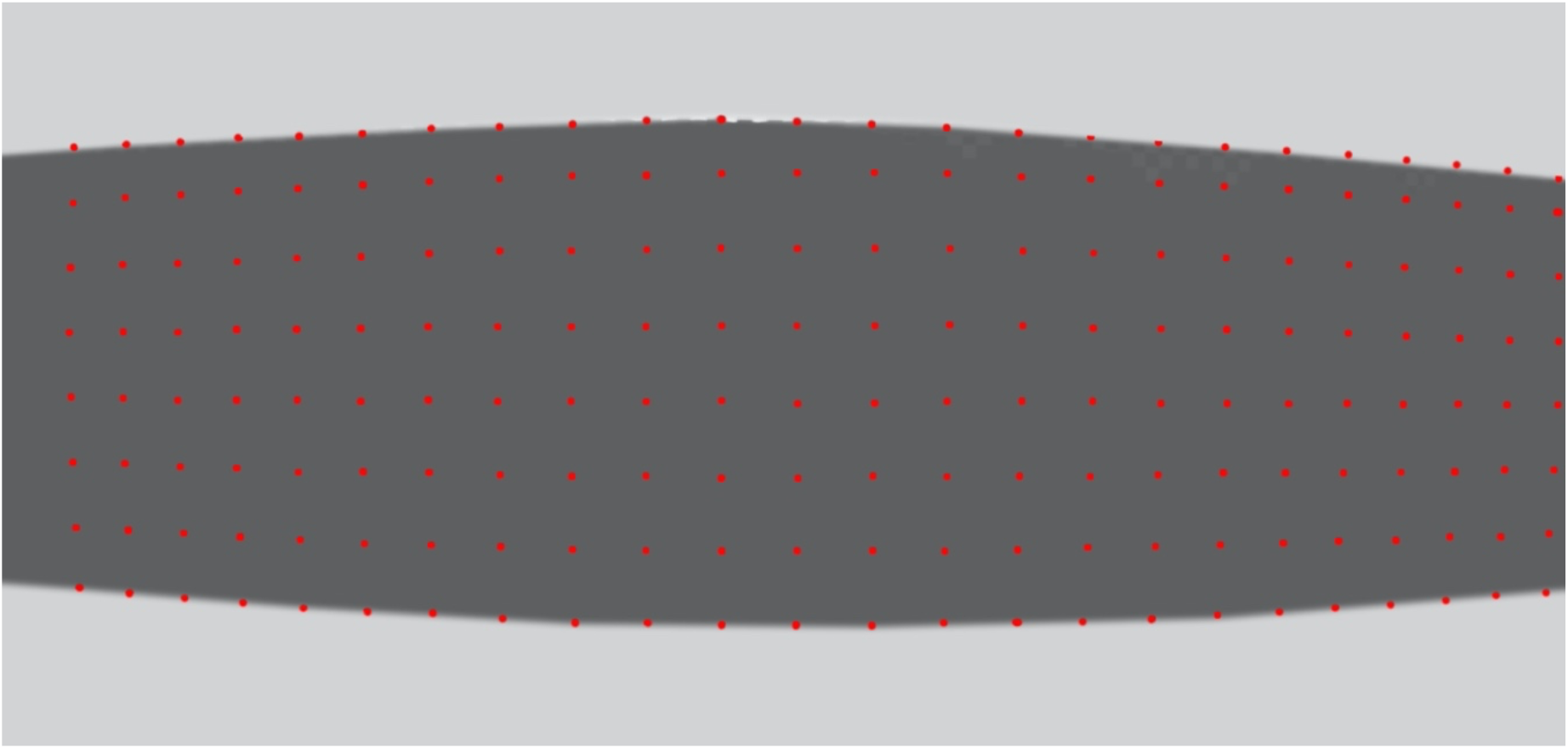
Camera view of the calibration grid on the floor (red points).

The intersection points of the grid in the camera image are digitised (manually, or by using specially developed image analysis software), and their pixel locations are recorded. Thus, each grid location (Xi,Yi) on the floor is tagged with its corresponding pixel co-ordinates (pxi,pyi) in the image. This data is used to compute a function that characterises a two-dimensional mapping between the grid locations on the floor and their corresponding pixel co-ordinates in the image. We note that the calibration grid does not need to be on the floor: It can be at any distance from the camera, but this distance must be known, and the plane of the grid must be perpendicular to the camera’s optic axis.

Video footage of a bird flying in the chamber, as captured by the overhead camera, is then analysed to reconstruct the bird’s 3D flight trajectory, as described below. Two examples of such footage are provided in the Supplementary videos SV1 and SV2. The positions of the wingtips are digitised in every frame in which the wings are fully extended, i.e. when the distance between the wingtips attains its maximum value (equal to the wingspan) in the video image. In the Budgerigar this occurs once during each wingbeat cycle, roughly halfway through the downstroke. We call these frames *Wex* frames, and denote the pixel co-ordinates of the wingtips in these frames by (pxL,pyL) (left wingtip) and (pxR,pyR) (right wingtip). The projected locations of the two wingtips on the floor are determined by using the mapping function, illustrated in Figure 2, to carry out an interpolation. Essentially, the projected location of this wingtip on the floor is obtained by computing the position of the point on the floor that has the same location, relative to its four surrounding grid points, as does the position of the wingtip (in image pixel co-ordinates) in relation to the positions of the four surrounding grid locations (in image pixel co-ordinates). Thus, in the case of the left wing tip, for example, this computation effectively uses the locations of the four grid points 1,2, 3 and 4 (see Figure 1) with locations (X1,Y1), (X2,Y2), (X3,Y3) and (X4,Y4) on the floor, and their corresponding image pixel co-ordinates (px1,py1), (px2,py2), (px3,py3) and (px4,py4) respectively, to interpolate the projected position of the pixel co-ordinate (pxL,pyL) on the floor. A similar procedure is used to project the position of the right wingtip (pxR,pyR) on the floor. The construction of the twodimensional mapping function, and the interpolation are accomplished by using the Matlab function *TriScatteredInterp*. (Equivalent customized codes could be written in any language.)

Once the positions of the two wingtips have been projected on to the floor, this information can be used to determine the instantaneous position of the bird in three dimensions, as illustrated in Figure 3. In this figure the 3D positions of the left and right wingtips are denoted by M, with co-ordinates (xL,yL,z), and N, with co-ordinates (xR,yR,z), respectively. Their projected points on the floor are denoted by C, with co-ordinates (XL,YL,0), and D, with co-ordinates (XR,YR,0), respectively.

**Figure 3.**
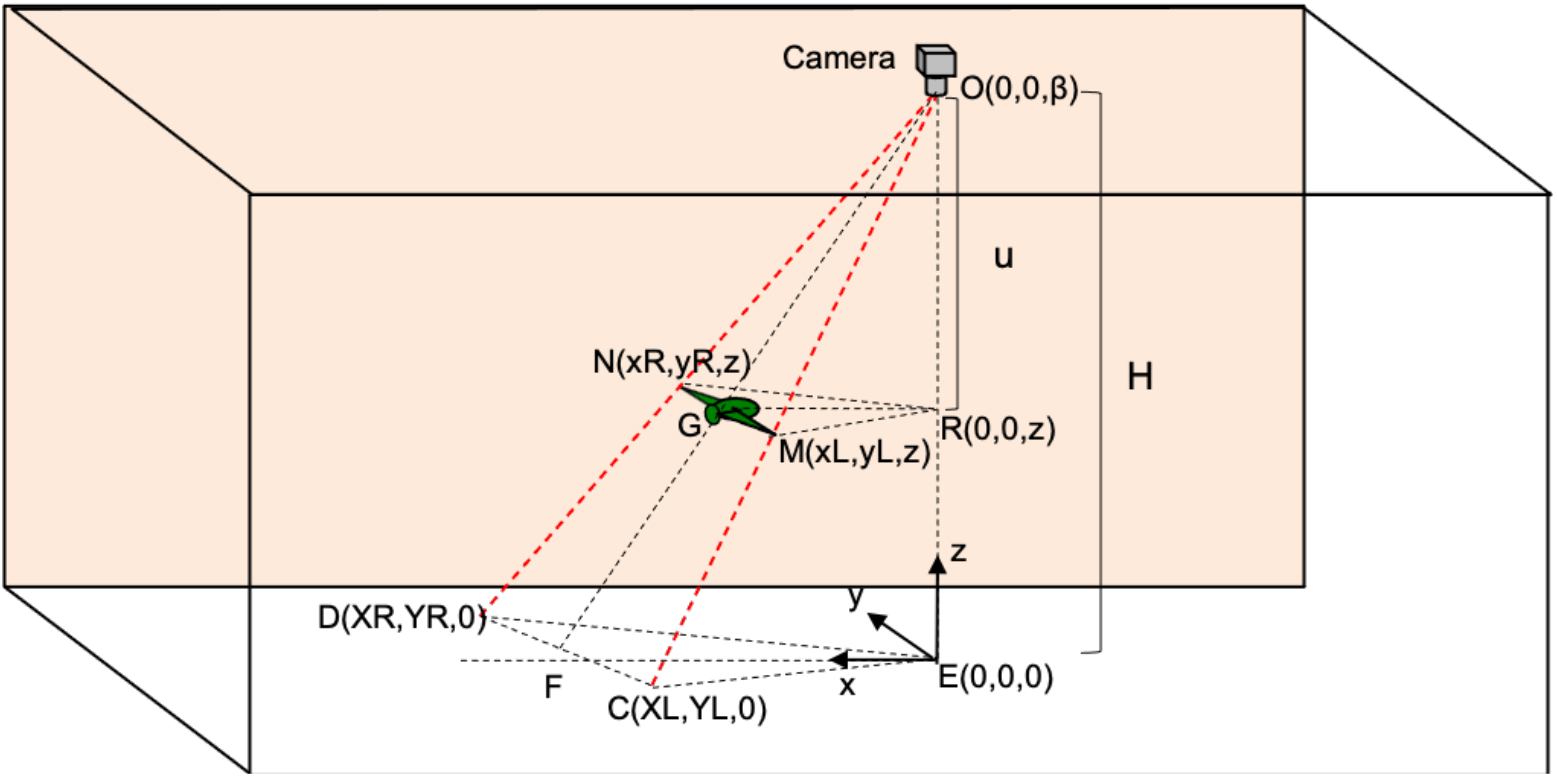
Schematic view of experimental chamber, showing the variables used for computing the instantaneous 3D position of the bird and its wingtips. E is the point on the floor that is directly beneath the camera, i.e. the point where the camera’s optic axis intersects the floor.

The height of the bird above the floor is established by determining the ratio between the known wingspan of the bird (w), and the projection of its wingspan on the floor, which we denote by W. W, which is equal to the distance between points C and D in Figure 3, is given by

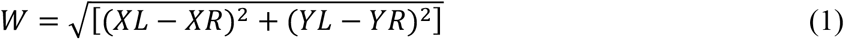

We denote the ratio (W/w) by Q.

From the geometrical similarity of the triangles OCD and OMN, and triangles OEF and ORG, we can write

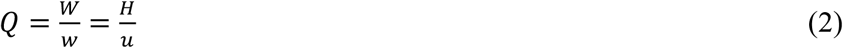

where H is the height of the ceiling (assumed to be known), and *u* is the distance of the bird below the ceiling. The height *h* of the bird above the floor, equal to (H-u), is then computed from (2) as

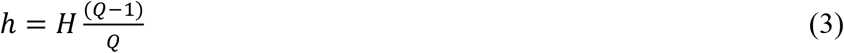

*h* is the height of the two wingtips above the floor. The (x,y) co-ordinates of the left and right wingtips can also be computed from the wingspan ratio Q as follows.

From the similarity of triangles ODF and ONG, and OEF and ORG, we have:

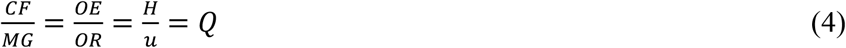

which can be rewritten as

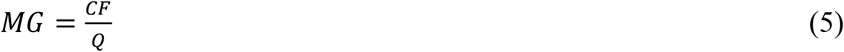

This implies that the (x,y) position co-ordinates of the left wingtip are given by

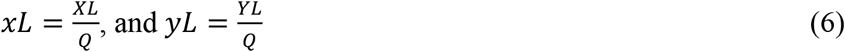

and the (x,y) position co-ordinates of the right wingtip are

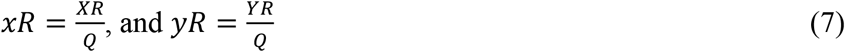

Thus, the 3D position co-ordinates of the left and right wing tips are (*xL,yL,h*) and (*xR,yR,h*). If we assume that the centre of the bird (the approximate position of its centre of gravity) is located midway between the extended wingtips (i.e. approximately at the thorax), then the 3D co-ordinates of the centre of the bird (*xc,yc,zc*) (which we shall henceforth refer to as the thorax) can be computed as

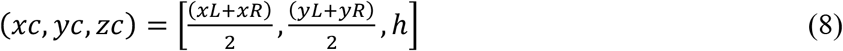

However, computing the centre of the bird in this way is valid only at the instants when the wings are fully extended. At other times the wings would be pointing either forward or backward, and this calculation would yield an incorrect result. Another approach would be to track the position of the head. During flight, the head is the most stable part of the bird’s anatomy-it maintains a horizontal orientation that is largely independent of the pitch and roll attitude of the body (Warrick et al., 2002; Frost, 2009; Bhagavatula, 2011). It is also a highly visible part of the bird that can be tracked reliably - either manually through frame-by-frame digitisation, or by software algorithms that employ relatively simple heuristics. Moreover, the head carries the bird’s primary sense organs, including the eyes. Therefore, reconstructing the 3D trajectory of the head can be useful for determining the visual stimuli that the bird experiences during its flight.

The 3D position co-ordinates of the head can be calculated for each frame as follows. The pixel co-ordinates of the head are determined in every frame (either through manual digitisation or an automated tracking algorithm). The head pixel co-ordinates are projected on to the floor, using the same interpolation procedure that was applied to the wingtips. We denote the floor co-ordinates of the head by (XH,YH) (not shown in Figure 3). Then, using the same geometrical reasoning as above, the (x,y) position co-ordinates of the head are given by

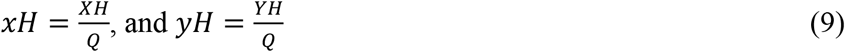

and the full 3D co-ordinates of the head are given by

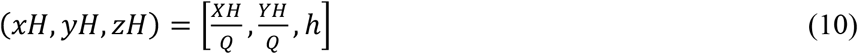

We note that the height of the head (*h*) is directly calculable only in the frames in which the wings are fully extended, because the bird’s wingspan is the known metric that enables determination of the height. The heights in other frames are estimated through temporal interpolation, assuming that the height varies approximately linearly between successive wing extensions. This is a reasonable assumption for most birds - typically, the height of flight varies slowly and smoothly across several wing beat cycles. However, the X and Y coordinates of the bird in 3D (*xH,yH*) are determined independently for each frame of the video sequence, based on the digitised pixel co-ordinates of the head in each frame, and the temporally interpolated height for that frame. Thus, while the height of the head (h) is temporally interpolated between wing extensions, the (X,Y) co-ordinates of the head (*xH,yH*) can be calculated independently for each frame, based on the pixel co-ordinates of the head in that frame. The height of the thorax can be estimated through a similar calculation, by defining the thorax either as the midpoint between the extended wingtips in the image, or the midpoint between the wing bases in the image.

In summary, our method delivers a sample of the bird’s height at every frame in which the wings are extended. These samples are interpolated in time to obtain a height profile of the head for the entire video sequence. This height profile is then used in combination with the pixel co-ordinates of the head in each frame to obtain the 3D co-ordinates of the head for each frame of the video sequence. The 3D trajectory of the thorax (defined as the mid-point between the extended wingtips) can also be reconstructed as described above.

In Budgerigars, the wings are fully outstretched only once during each wingbeat cycle –roughly halfway through the downstroke, as we have noted above. This also appears to be the case in pigeons and magpies (Tobalske and Dial, 1996). It is possible that in certain other species, which move their wings in the same plane during the upstroke and the downstroke, without folding them, there are two *Wex* frames per wingbeat cycle - one occurring during the upstroke, the other during the downstroke. In such cases we can obtain two height estimates per wingbeat cycle, and therefore reconstruct the height profile at twice the temporal resolution.

In the above analysis, we have assumed that the head of the bird is at the same height as that of its extended wingtips. If the head is at a different height - as may be evinced from prior knowledge or from side-view images of bird flight in wind tunnels - this known height offset can be added to the wingtip height to obtain the true height of the head.

### 2.2 Procedural steps

Based on the theory described above, the step-by-step procedure for reconstructing the 3D trajectory of the head of a bird from a video sequence captured by a single overhead camera can be described as follows:

i. Construct the floor grid and acquire an image of the grid from the video camera. An example is shown in Figure 2. The grid is used only once for the camera calibration, and does not need to be present in the experiments.
ii. Digitise the pixel co-ordinates of the grid locations in the camera image, to obtain a one-to-one mapping between the real co-ordinates of the grid locations on the floor and their corresponding pixel coordinates in the image.
iii. Acquire knowledge of the bird’s wingspan, either from published data for the species, or, preferably, from direct measurement of the actual individual (because the wingspan can vary slightly across individuals due to age and other factors).
iv. Acquire video footage of the bird during flight in the chamber
v. Select the frames in the video sequence in which the wings are fully extended. The selection can be done either manually, or through custom-written software. The wing-extension frames are denoted by *Wex*.
vi. Digitise the pixel positions of the left and right wingtips of the bird in each of the *Wex* frames, as shown in the illustrative example of Figure 4.
vii. Determine the height of the thorax (midpoint between the extended wingtips) or of the head in each of the *Wex* frames from equations (1-3).
viii. Obtain the height profile of the thorax (and/or the head) for the entire video sequence by temporally interpolating the heights calculated for the *Wex* frames.
ix. Digitise the pixel position of the thorax/head in each frame of the video sequence.
x. Compute the 3D position of the thorax/head for each frame from equations (9) and (10).

**Figure 4.**
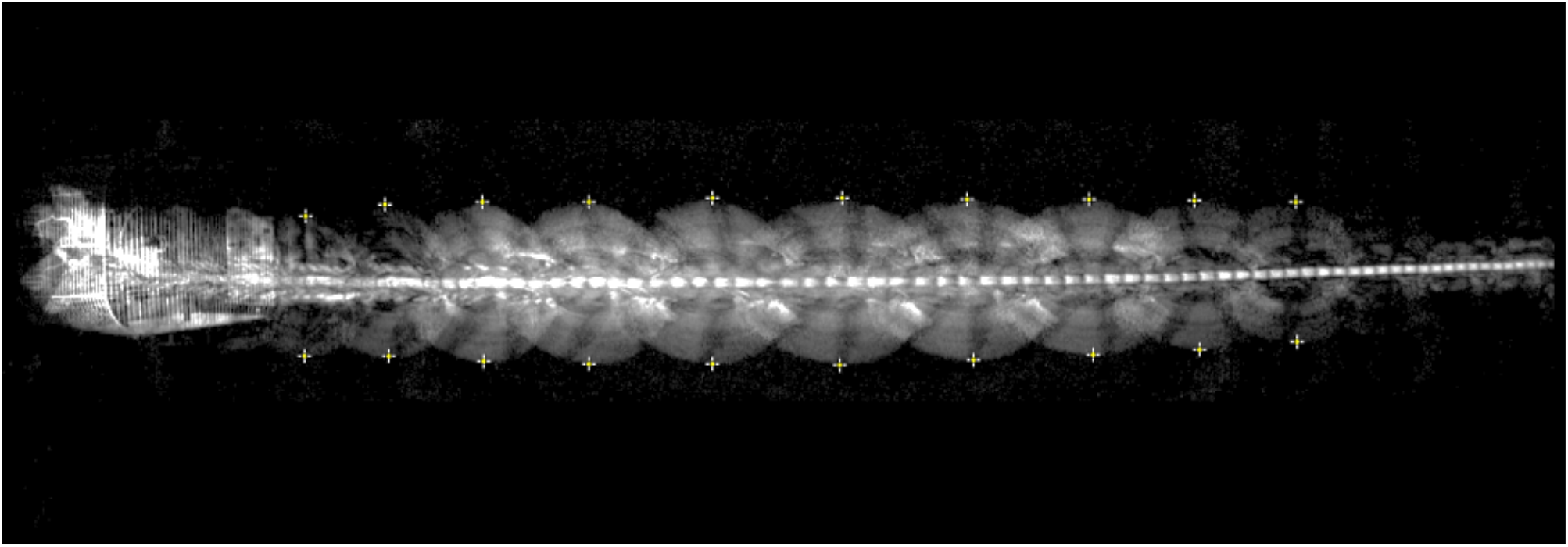
Example of a video sequence showing superimposed images of the bird in successive frames. Successive wing extensions are marked by the crosses.

### 2.3 Test of accuracy

The precision of the 3D trajectory reconstruction procedure was evaluated by placing a small test target at 44 different, known 3D locations within the tunnel, of which 39 were within the boundary of the grid. The test target was a model bird with a calibrated wingspan of 30 cm. The head was assumed to be midway between the wingtips, and at the same height as the wingtips. This assumption does not affect the generality of the results, as discussed above. The standard deviations of the errors along the x, y and H directions were 2.1 cm (X), 0.6 cm (Y) and 2.6 cm (H). A detailed compilation of the errors is given in Table S1 of the SI.

### 2.4 Ethics Statement

All experiments were carried out in accordance with the Australian Law on the protection and welfare of laboratory animals and the approval of the Animal Experimentation Ethics Committees of the University of Queensland, Brisbane, Australia (Animal Ethics Approval Certificate QBI/075/18).

## 3. Results

### 3.1 Examples of flight tracking and reconstruction

Here we show some examples of reconstruction of 3D trajectories of flights of Budgerigars through an indoor tunnel, of dimensions of dimensions 7.28 m (length) x 1.36 m (width) x 2.44 m (height). The birds were trained to fly from a perch at one end of the tunnel to a bird cage at the other. A downward-facing video camera, placed at the centre of the ceiling of the tunnel, was used to film the flights and reconstruct the trajectories in 3D. A grid, of check size 20 cm x 20 cm (as in Figure 2), was drawn on the floor to calibrate the camera using the procedure described above. The reconstructed 3D trajectories do not include the take-off and landing phases of the flight. They only show a section of the trajectory within a window that extends from about 1.75 m ahead of the camera aperture to about 0.25 m behind it, which could be regarded as a ‘cruise’ phase where the bird has completed take-off and not yet commenced landing.

Flights through the tunnel were filmed with the tunnel being either empty (devoid of any obstacles) or carrying a narrow, vertically oriented aperture (a slit) at the halfway point, through which the birds had to fly to get to the other end. To prevent injuries to the birds, the aperture was created by suspending two cloth panels that reached from the ceiling to the floor. Two aperture widths were tested: In one set of tests, the aperture was 5cm wider than the bird’s wingspan; in the other set, the aperture was 5cm narrower than the bird’s wingspan. It has been shown in earlier studies (Schiffner et al., 2014; Vo et al., 2016) that Budgerigars are acutely aware of their wingspan: when negotiating a narrow aperture, they fold their wings back briefly only when the aperture is narrower than their wingspan, and fly through without interrupting their wingbeat cycle if the aperture is wider than their wingspan.

A plan view of a reconstructed flight is shown in Figure 5. In this example, the bird (*Four*) has a wingspan of 29 cm and it flies through a 34 cm aperture, which is 5 cm wider than the wingspan. The figure shows the (X,Y) positions of the two wingtips at the time of each wing extension, the yjorax (defined as the midpoint of the line joining the extended wingtips), and the position of the head. It is evident that the bird flies through the aperture without interrupting its wingbeat cycle, as the wingbeat extensions are equally spaced.

**Figure 5.**
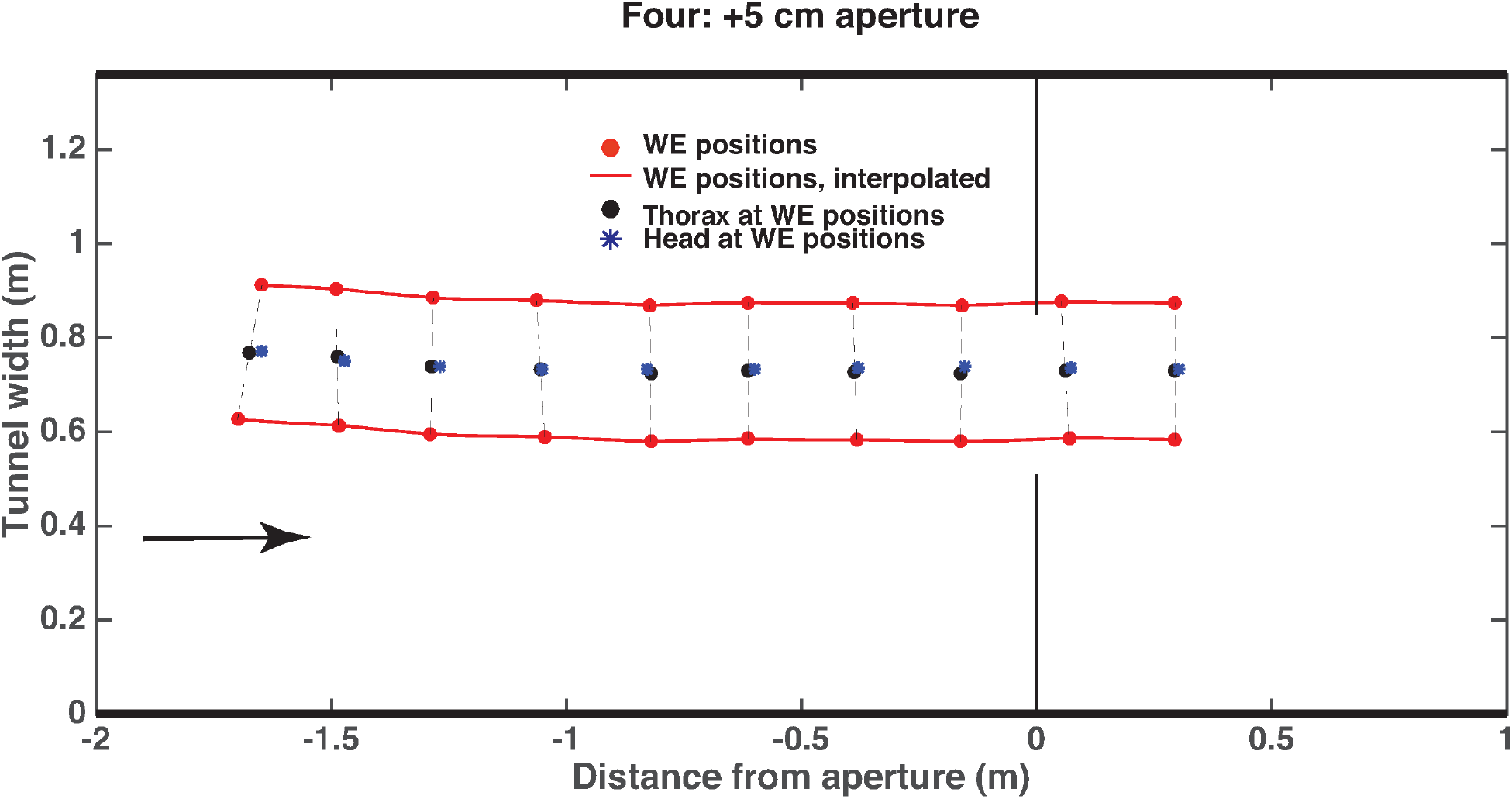
Plan view of a reconstructed flight of Bird Four. In this example, the wingspan of the bird is 29 cm and it flies through a 34 cm aperture, which is 5 cm wider than the wingspan. Details in text. The red circles show the wingtip positions at the time of each wing extension, the black circles show the inferred position of the thorax at these instants, and the blue asterisks depict the position of the head at these instants. The red lines show the wing extension trajectories interpolated between wing extensions. The arrow in this and other figures shows the direction of flight.

This is also clear from Figure 6, which shows two 3D views of the same flight trajectory, where the blue circles represent the position of the thorax at each wing extension and the red curve shows the reconstructed 3D position of the head for every frame, as described in the text above and in the legend. The lateral view of the trajectory (Figure 5, right hand panel) shows that the bird maintains its height while passing through the aperture, because the wingbeat cycle is not interrupted.

**Figure 6.**
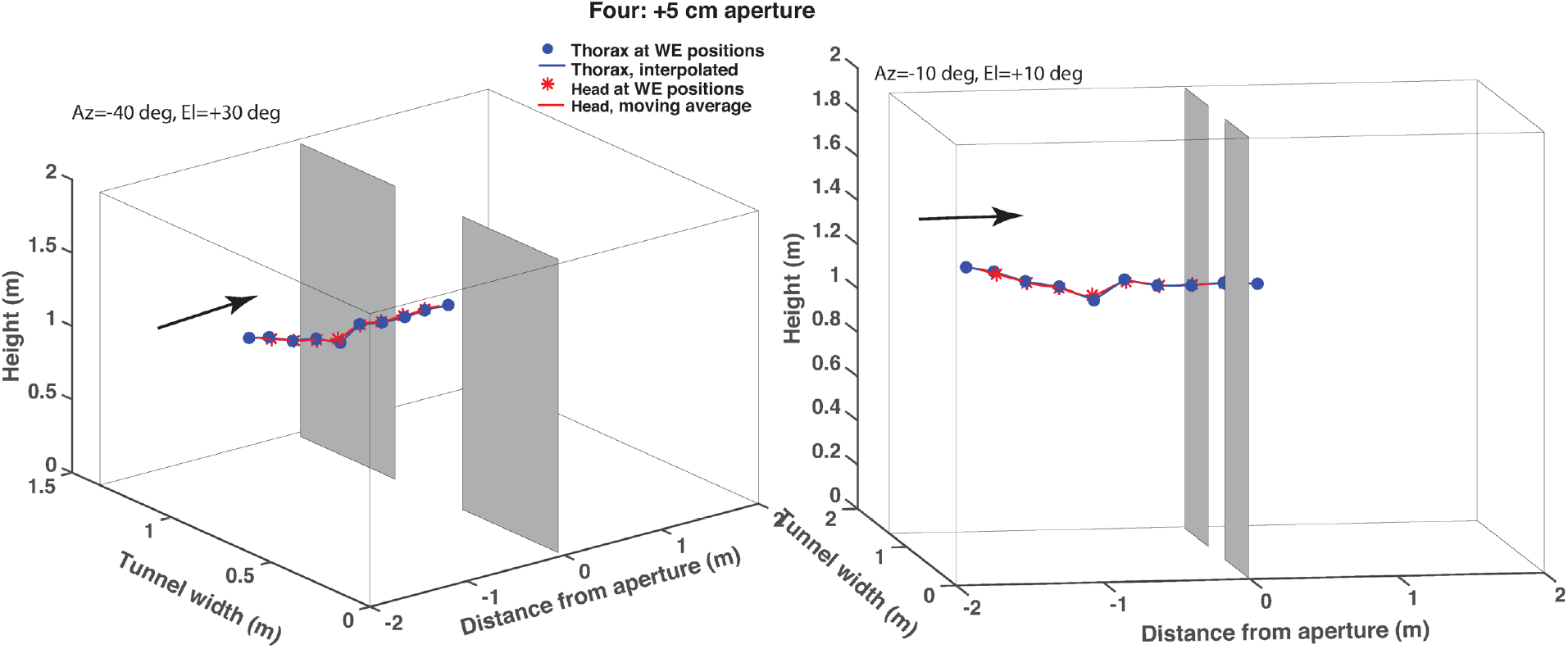
Two 3D views of the trajectory shown in Figure 5, in which bird Four flies through an aperture that is 5 cm wider than its wingspan. The blue circles show the inferred position of the thorax at the time of each wing extension, the blue lines show the linearly interpolated thorax positions between successive wing extensions, and the red asterisks show the head position at the time of each wing extension. The image coordinates of the head, which were digitized in every video frame, were used to calculate the 3D trajectory of the head in every frame, as described in the text. The red curve shows the resulting 3D trajectory of the head during the entire video sequence, after smoothing by a 9-point centered rectangular moving average filter. Left panel: View from −40 deg azimuth, 30 deg elevation. Right panel: Near-lateral view from −10 deg azimuth, 10 deg elevation.

Figure 7 shows two 3D views of a trajectory of the same bird during flight through an aperture that is 5 cm narrower than its wingspan. Here is clear that the wingbeat cycle is interrupted when the bird passes through the aperture - the distance between successive wing extensions is dramatically larger during the passage. This is also clear from the lateral view of the trajectory (Figure 6, right hand panel), which shows that the bird loses altitude while passing through the aperture, because the wingbeat cycle is interrupted.

**Figure 7.**
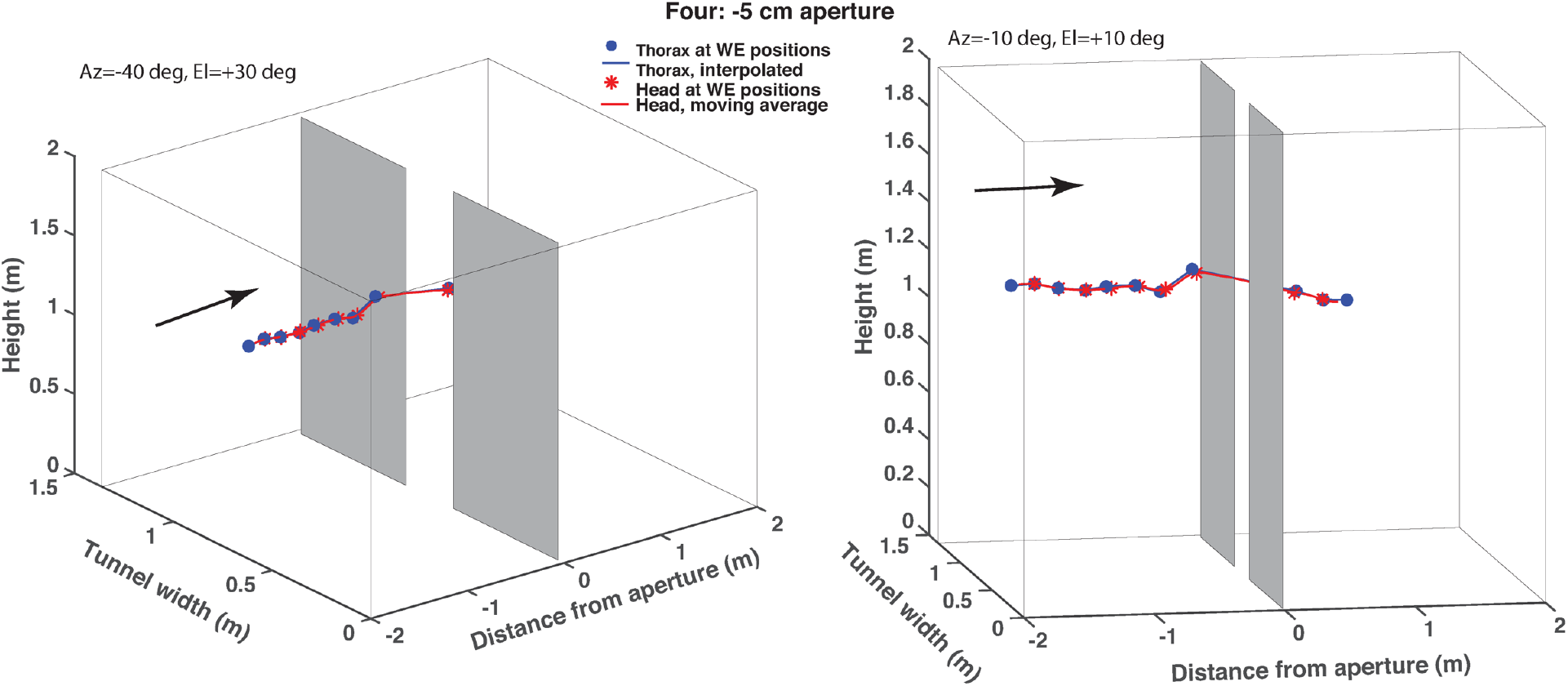
Two 3D views of a trajectory of bird Four during flight through an aperture that is 5 cm narrower than its wingspan. Details are as in Figure 6.

Figure 8 shows two 3D views of a trajectory of the same bird during flight through the tunnel when there is no aperture. In this case – as in Figure 5 - the wingbeat cycle is not interrupted anywhere in the flight. This is also clear from the lateral view of the trajectory (Figure 7, right hand panel), which shows that the bird maintains a constant wingbeat cycle and does not lose altitude abruptly anywhere along the trajectory.

**Figure 8.**
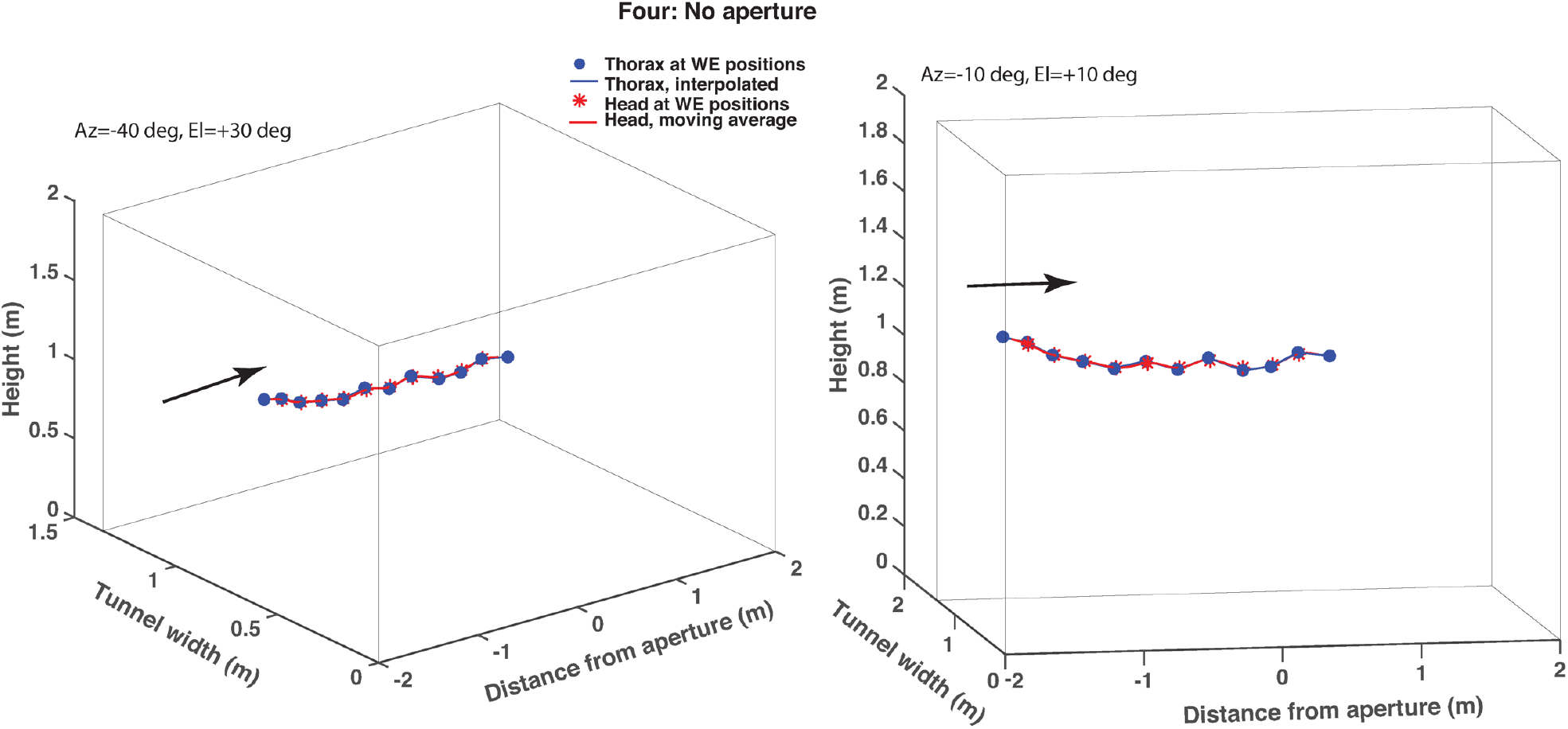
Two 3D views of a trajectory of bird Four during flight through a tunnel which carries no aperture. Details are as in Figure 6.

It is clear from Figs. 5 – 8 that bird *Four* interrupts its wingbeat cycle only when it confronts an aperture that is narrower than its wingspan, and not when the aperture is wider than the wingspan or is not present in the tunnel. A loss of altitude occurs only when the wingbeat cycle in interrupted, and not otherwise.

Figure 9 shows plan views of the reconstructed 3D trajectories of the head for the three conditions. In each case, the asterisks mark the locations of the head at the times of full wing extension. Other details are given in the figure legend. In the case of the narrow aperture (red track), the bird temporarily interrupts its wing beat cycle while passing through the aperture. The final wing extension prior to passing the aperture occurs at a point approximately 0.35 m ahead of the aperture. The wing beat cycle resumes after passage through the aperture, with the first wing extension occurring at a point approximately 0.5m beyond the aperture. In the wide aperture and the no aperture conditions, the wing beat cycle continues uninterrupted throughout the flight. These observations are in agreement with those of Schiffner et al. (2014), who report an exquisite ability of these birds to gauge the width of oncoming passages in relation to their wingspan. However, their study only recorded the frequency and timing of wing closures, and did not reconstruct the birds’ trajectories in 3D.

**Figure 9.**
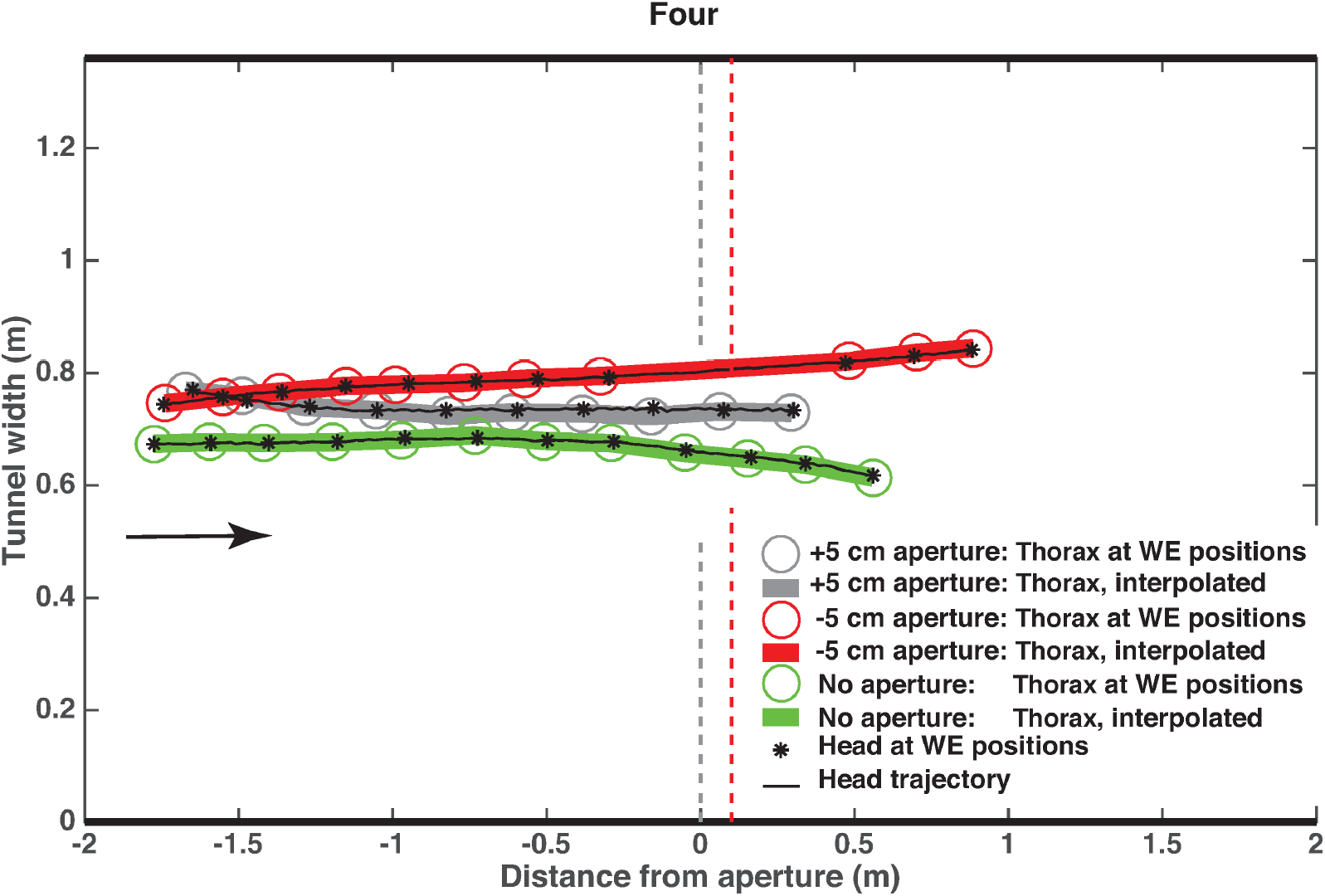
Plan views of the reconstructed 3D trajectories for the narrow aperture condition (red), the wide aperture condition (grey) and the no aperture condition (green), for bird Four. In each case, the circles mark the locations of the thorax (defined as the mid-point of the line connecting the extended wing tips) at the time of each wing extension, the thick curves show the thorax trajectory interpolated from the thorax positions at these times, the asterisks mark the locations of the head at the times of the wing extensions, and the thin black curve through the asterisks shows the trajectory of the head, reconstructed from the digitized image co-ordinates of the head in each frame as explained in the text.

Figure 10 shows reconstructed profiles of the forward flight speed (speed along the X axis of the tunnel) for the flights of bird *Four* in the narrow aperture, wide aperture and no-aperture conditions. These profiles were constructed using three different procedures, the details of which are described in the legend. The three procedures yield consistent results. The principal observation is that the forward speed is more or less constant throughout the flight and is independent of the flight condition, as observed in Vo et al. (2016). Interestingly, the interruption of the wing beat cycle during the flight through the narrow aperture does not significantly reduce the forward speed.

**Figure 10.**
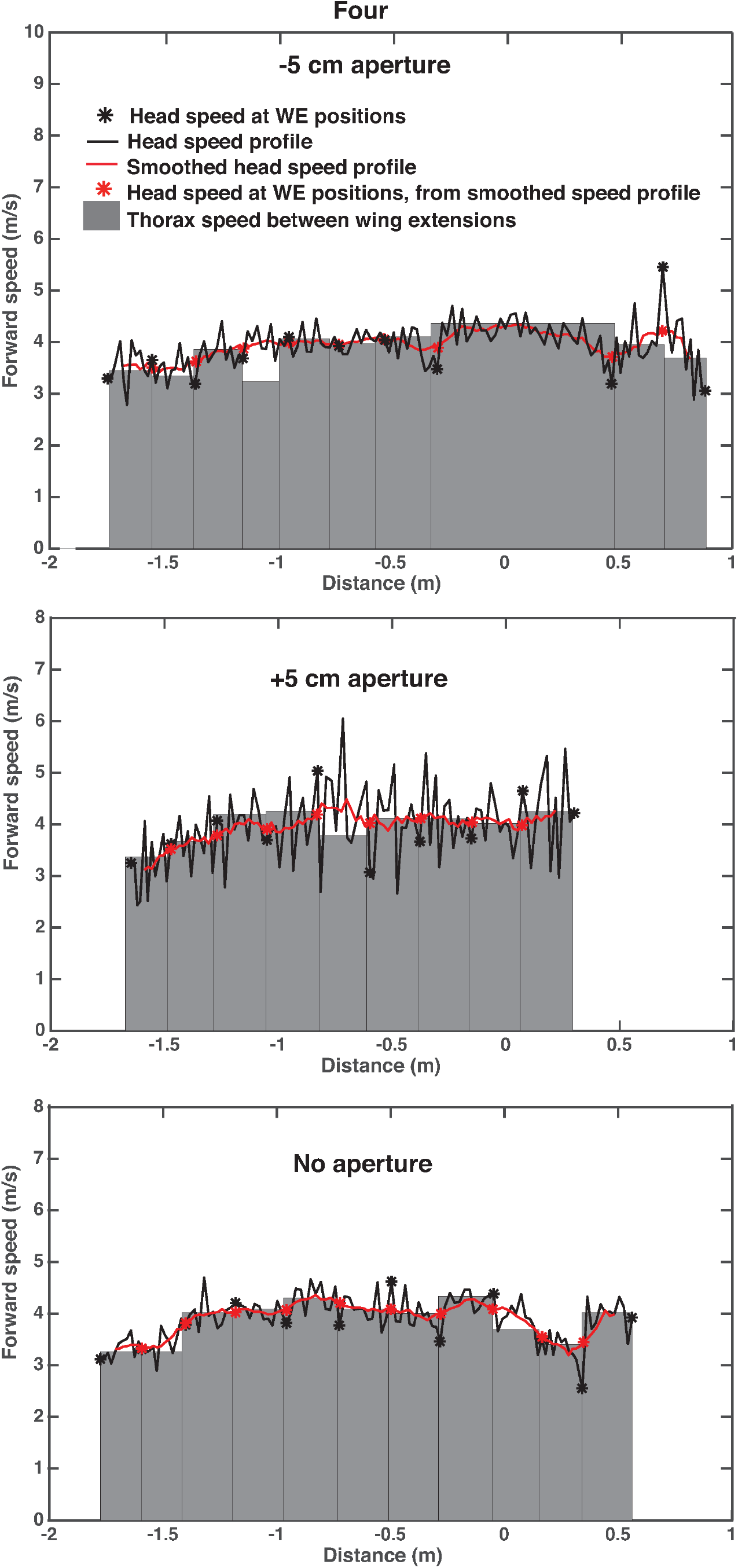
Forward speed profile of bird Four during flight through the narrow aperture (top panel), the wide aperture (middle panel), and the empty tunnel (bottom panel). In each case, the black curve shows the speed profile of the head, computed from the frame-to-frame X positions of the head. The black asterisks denote the speeds at the head positions corresponding to the wing extensions, and the red curve and asterisks depict the result of smoothing the speed profile using a 9-point centered rectangular moving average filter. The edges of the grey bars denote the successive X positions of the thorax (defined as the midpoint between the wingtips) at each wing extension, computed as described in the text. The height of each grey bar depicts the mean forward speed of the thorax between successive wing extensions, computed as the ratio of the X distance between successive edges, to the time interval between these edges. Because the wingbeat kinematics are not perfectly identical from one wingbeat cycle to the next, and the head could make small movements relative to the thorax, the spatial relationship between the head and the line joining the wingtips is not exactly the same at the point of each wing extension. As a result, the measured speed of the thorax (grey bars) can occasionally be noticeably different from that of the head (asterisks): e.g. fourth grey bar (top panel), and fifth grey bar (middle panel).

In the Supplementary Information (Figures S1-S6) we show results for another bird (*Nemo),* corresponding those shown above for bird *Four.*

### 3.2 Accounting for the effects of body roll: extended calculation

In the analysis so far, we have assumed that the birds are not rolling during flight, i.e. that the wingtips are in the horizontal plane when they are fully extended. This assumption is quite valid for the flights we have filmed and reconstructed: the birds displayed very little roll throughout their flight, as evinced by the fact that, when the wings were outstretched, the two wingtips were approximately equidistant from the head in all video frames. However, our theory can be extended to take body roll into account – when this is significant – and continue to obtain accurate estimates of the 3D trajectories, as well as the roll angles. The essential elements of the procedure are described briefly below, and the complete general derivation is provided in the Supplementary Information (Section B).

The basic principle underlying the calculation of the roll angle is outlined in Figure 11, which illustrates a simplified case in which the midpoint O between the extended wingtips is directly under the camera, i.e. in line with the camera’s optic axis. During a roll, the line connecting the extended wingtips is not horizontal. Therefore, in the camera image the angles ϕ_1_ and ϕ_2_ subtended by the two wings will not be equal, because the right wingtip (in this case) is higher than the left wingtip. These angles can be computed from the projections on the grid floor of the images of the two wingtips (R,L), and the projection of the point (O), which is the point on the bird that is midway between the extended wingtips. When the bird is not rolling the extended wingtips will lie in a horizontal plane, and the image of O in the camera will be midway between the images of the extended wingtips, because ϕ_1_ = ϕ_2_. However, when the bird is rolling, ϕ_1_ ≠ ϕ_2_ and the image of O will not be midway between the images of the extended wingtips. ϕ1 and ϕ2 can be measured and used to calculate the roll angle. In the camera image, O is determined as the point where the straight line connecting the wingtips intersects the longitudinal axis of the thorax (see Figure 13c). We shall hereafter refer to O as the ‘thorax point’. From the projected locations of R, L and O on the floor grid, the height of O above the ground (h) and the angle of roll (a) can be calculated as shown below. This calculation distinguishes between changes in roll and changes in height, because it evaluates the distances of the projected wingtips *separately* for the left and right wings: these distances are not equal when the bird rolls.

**Figure 11.**
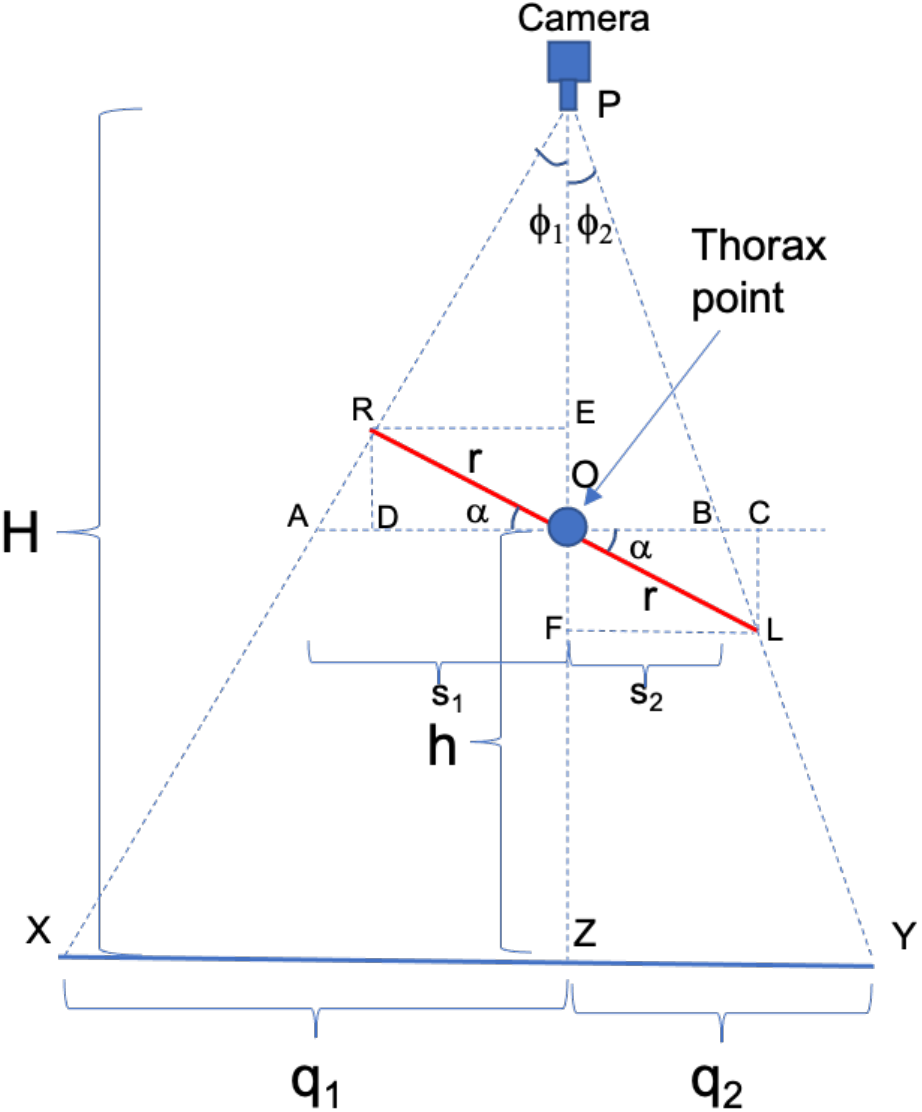
Illustration of calculation of bird height (h) and roll angle (a)for a simplified case in which the midpoint of the wingspan is directly below the camera’s optical axis. The known wingspan of the bird is 2r, and the camera is at a known height H above the floor.

From Figure 11 we have

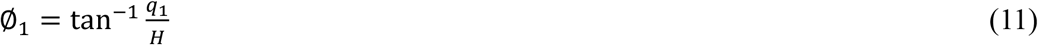

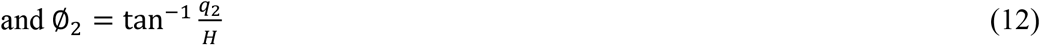

so that

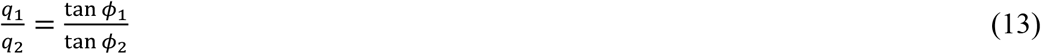

We can also write

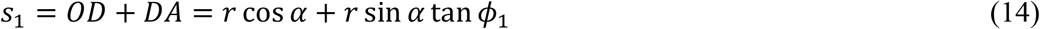

and

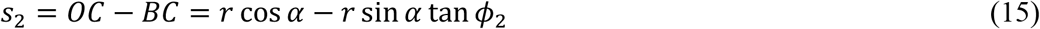

Thus,

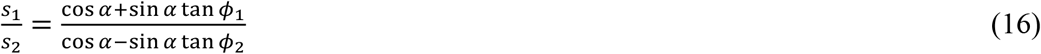

From triangle similarity we have

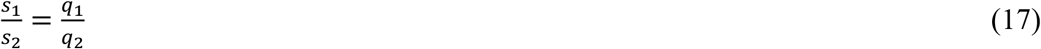

Equating (13) and (16) we obtain

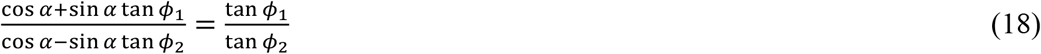

which gives

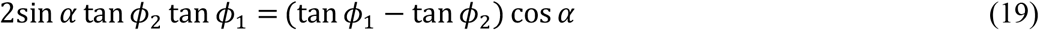

or

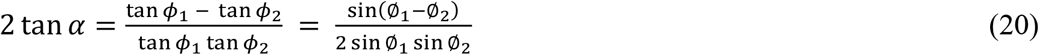

from which we obtain

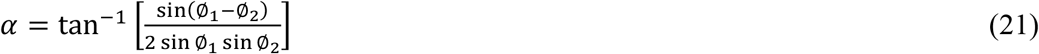

Thus, the roll angle a can be evaluated from (21), using (11) and (12) to calculate Ø_1_, and Ø_2_.

Now, considering the total length AB, we have

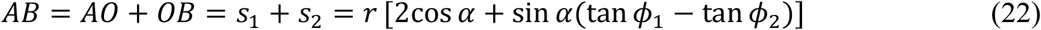

Then, from triangle similarity we have

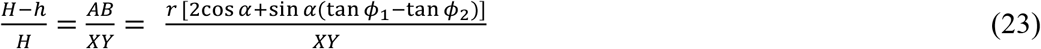

from which we solve for *h* to obtain

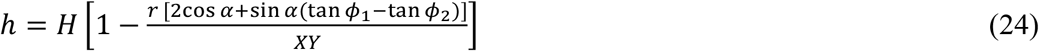

Since X and Y are the projections of the right and left wingtips on the floor, the distance [XY] can be calculated, and *h* can be evaluated using (24).

Note that Figure 11 illustrates a very simple case, for the purpose of conveying the basic approach to the calculation. In the Supplementary Information (Section B) we derive an extension of this calculation for a general case in which the bird can be at any 3D location. The extended calculation delivers the height of the bird, as well as the roll angle. Once the height is known the 3D coordinates of the head can computed as before, following the procedure described in Section 2.1 [equation (10)].

### 3.3 Validation of extended calculation

We have validated the extended calculation, which accounts for body roll, by using a model bird in a scaled-down arena with an overhead camera and a calibration grid on the floor. A calibrated platform was used to position the model bird at 20 different 3D locations and flight directions, at various known roll angles and heights above the floor (Figure 12).

**Figure 12.**
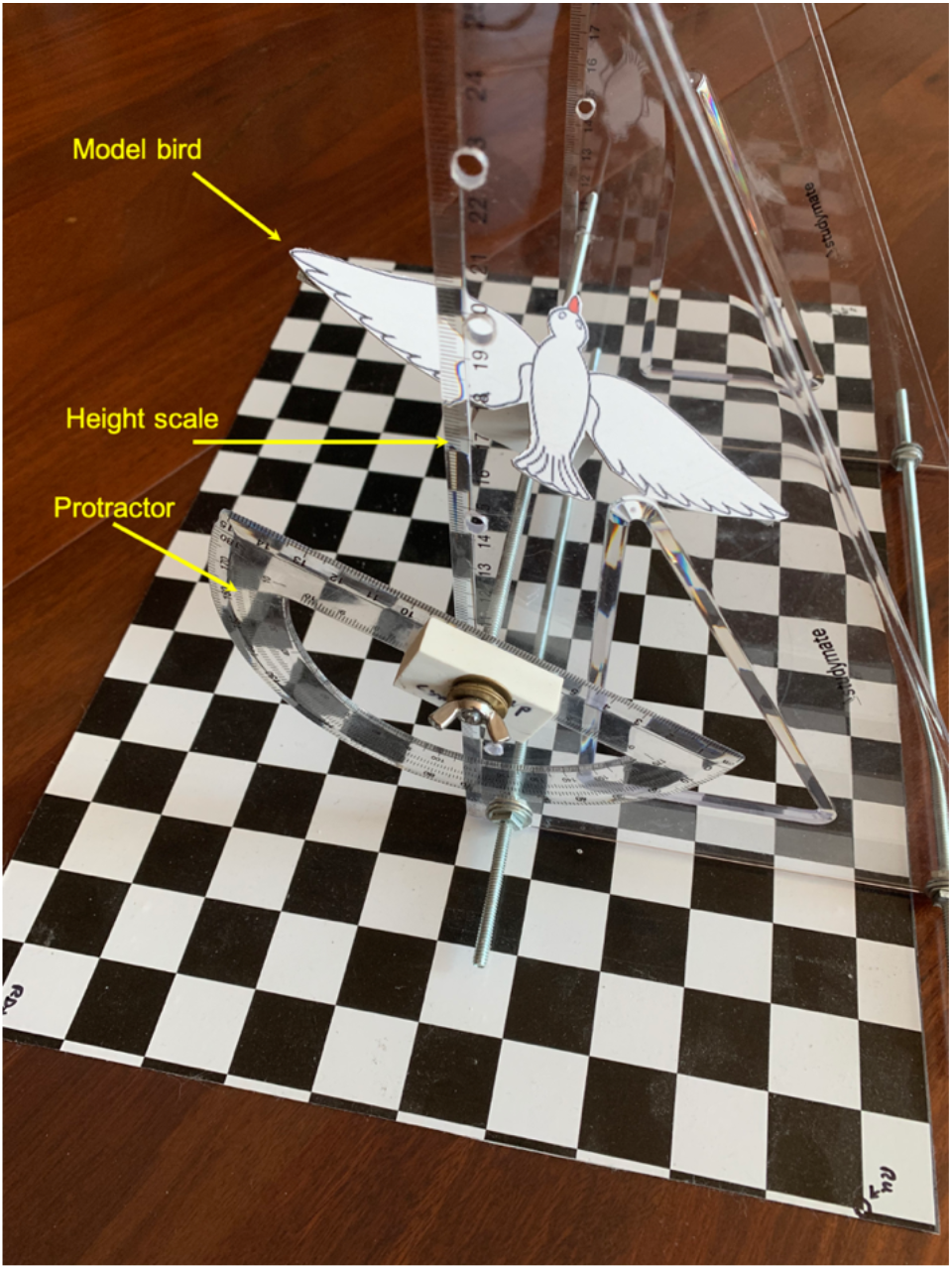
Setup for validation of extended calculation, which accounts for body roll. The model bird, with a wingspan of 180 mm, is mounted on a platform whose height and tilt (roll) can be set to calibrated values using the height scale and the protractor. The size of the floor grid is 408 x 260 mm, and the individual checks are 26.0 x 25.5 mm. The overhead camera is positioned above the center of the grid, with the nodal point of its lens at a height of 448 mm.

Four examples of the images captured by the overhead camera are shown in Figure 13.

**Figure 13.**
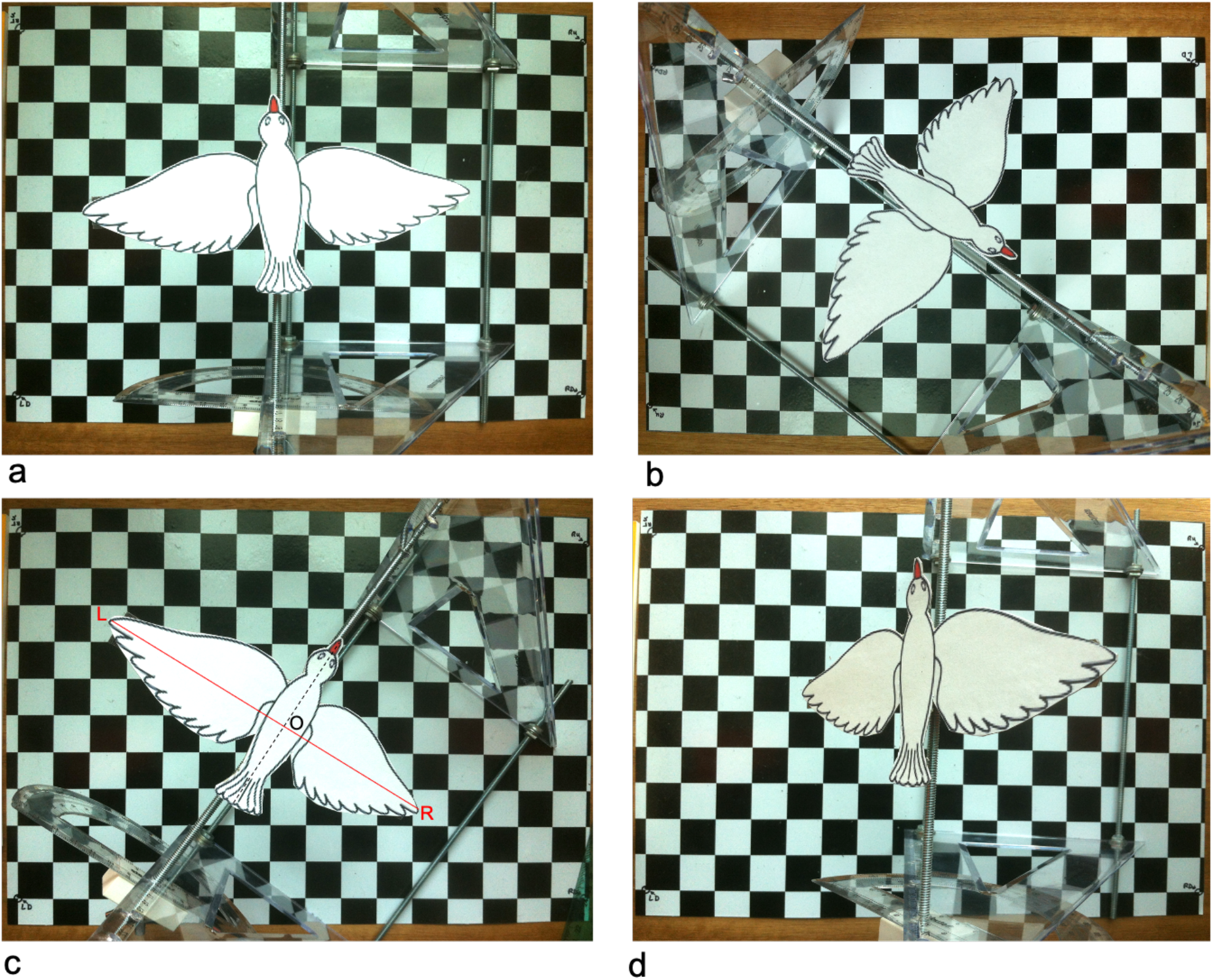
Examples of images acquired by the overhead camera. (a) Bird height 158mm, roll 0 deg; (b) Bird height 158mm, roll +26 deg; (c) Bird height 157mm, roll −22.5 deg; (d) Bird height 205 mm, roll +51 deg. Heights refer to the height of the thorax point O, as illustrated in (c), and in Figures 11, S7 and S8. The roll angle is positive when the right wingtip is higher than the left wingtip, and vice versa.

Figure 14 compares the thorax heights and roll angles derived from the extended calculation with the true (ground truth) values, for each of the 20 configurations. It is evident that the extended analysis delivers accurate estimates of the height of the bird irrespective of body roll, as well as accurate estimates of the roll angle.

**Figure 14.**
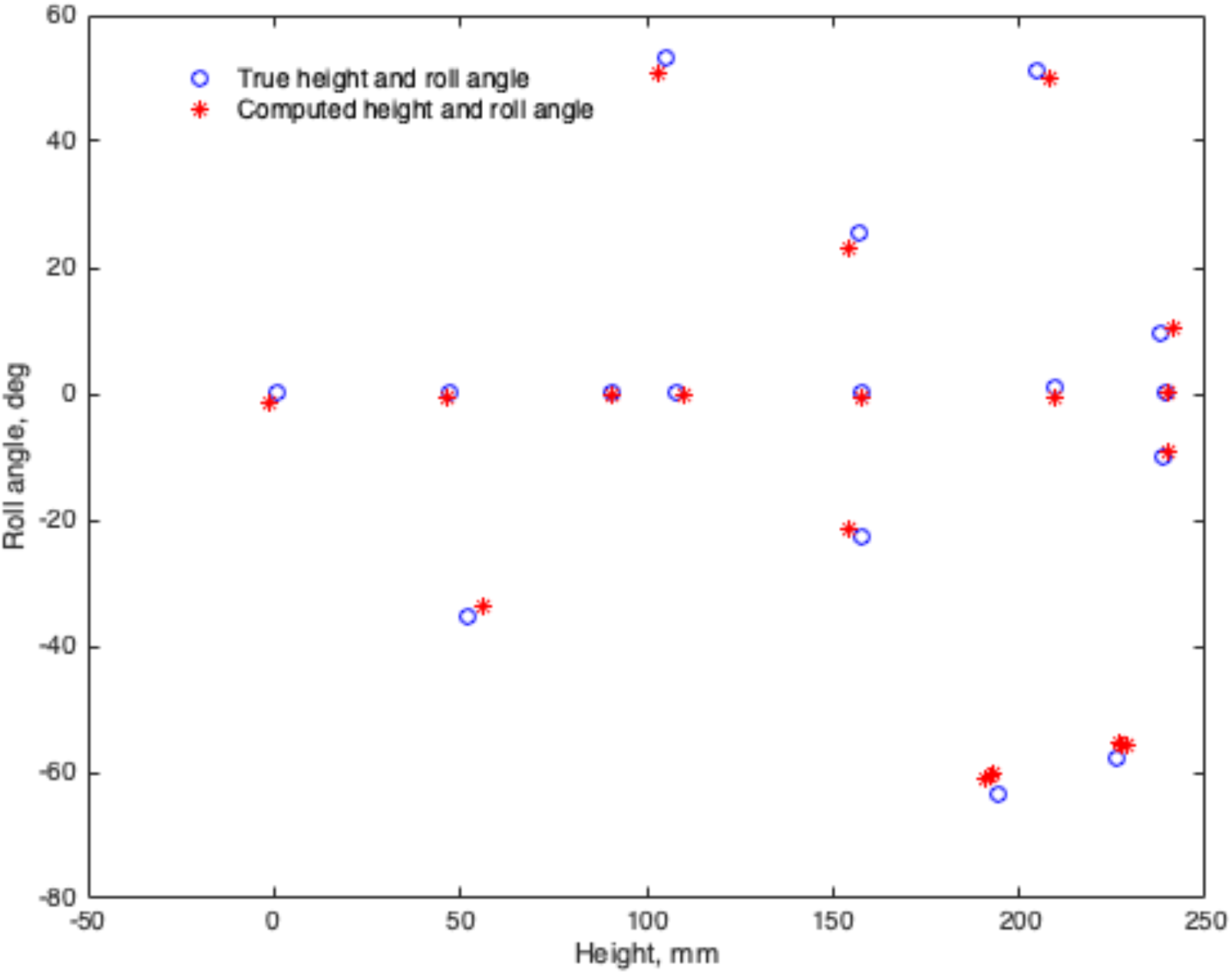
Examples of images acquired by the overhead camera. (a) Bird height 158mm, roll 0 deg; (b) Bird height 158mm, roll +26 deg; (c) Bird height 157mm, roll −22.5 deg; (d) Bird height 205 mm, roll +51 deg. Heights refer to the height of the thorax point O, as illustrated in (c), and in Figures 11, S7 and S8. The roll angle is positive when the right wingtip is higher than the left wingtip, and vice versa.

For the results shown in Figure 14, the mean and RMS errors in the calculated heights are 0.06 mm and 2.3 mm, respectively (0.03% and 1.3% of the wingspan, respectively), and the mean and RMS errors in the calculated roll angles are −0.4 deg and 1.7 deg, respectively.

While the real birds did not display significant body rolls during their flights in our experimental tunnel, the validation of our extended calculation using the model bird demonstrates that this procedure can be applied to calculate the true height (and the 3D coordinates of the thorax point and the head) of a bird even when its body is rolling, and to compute the roll angle.

## 4. Discussion

This study has described a simple, inexpensive method for reconstructing the flight trajectories of birds in 3D, using a single video camera. The advantages of the method are:

i. The technique does not use a conventional stereo-based approach. Therefore, it does not require complex calibration procedures involving capturing views of a checkerboard at various positions and orientations, which does not always guarantee accurate localisation in all regions of the experimental space.
ii. The technique does not need feature correspondences to be determined across video frames from two or more cameras.
iii. The grid marker on the floor provides a calibration of the camera geometry and accounts for all of the distortions in the camera optics. There is no need to assume that the camera can be approximated by a pinhole camera, or by any other specific optical geometry. This calibration is a one-off procedure that can be used for the rest of the lifetime of the camera/lens combination, provided the optics are not altered.
iv. Once the calibration has been performed, the calibration grid can be removed or covered (if this is necessary to prevent its potential influence on the behaviour of the birds in the experiments).
v. When a bird glides with its wings outstretched, its height (and therefore the 3D coordinates of the wingtips, the thorax point and the head) can be reconstructed in every frame without requiring any interpolation.
vi. Moving the camera to a different location does not require recalibration. 3D trajectories of birds can continue to be reconstructed with reference to the new optical axis of the camera and the new plane of the (internally stored) calibration grid. Thus, in principle, the camera that was calibrated in our experiments using the calibration grid on the floor, can also be used to reconstruct the trajectories of birds in outdoor flight by facing the camera upwards and performing the reconstruction relative to the same calibration grid. Trajectory reconstruction is possible even if a bird is located on the opposite side of the calibration grid – the geometry and interpolation underlying the reconstruction will be the same. This is a major attribute of the system, because - unlike systems that use stereo or multiple cameras – it does not need to be recalibrated every time it is moved to a new location.
vii. Because the method is computationally simple, it can be applied in closed-loop experimental paradigms in which visual stimuli need to be modified in real time in response to the bird’s flight, as is now being done with some animals (e.g. Stowers et al., 2017).
viii. The system is compact, portable, and easily deployed in the field.

The limitations of the method are:

i) We have assumed that the wings are fully extended at each extension, and that the tip-to-tip distance at these extensions is always equal to the measured wingspan. Variability in the wingtip distances from extension to extension (which may occur during certain manoeuvres) will introduce errors in the reconstruction of the 3D trajectory.
iii) The calibration grid on the floor grid must cover a sufficiently large area to enable projection of the wingtips on to the floor at all possible bird positions. This could be a problem when the bird is flying close to the ceiling or to one of the walls of the tunnel (or chamber), as it would require extrapolation of the grid beyond the floor of the chamber. Grid extrapolation can be carried out, but it requires assumptions to be made about the unknown optical distortions in the extrapolated regions of the grid. The calibration grid does not need to be on the floor: it can be in a parallel plane that is much closer to the camera – for example, a few centimetres away. In this case the grid can be considerably smaller, but it must be large enough to span the entire visual field of the camera.
iv) The method requires selection of the *Wex* frames in the video sequence, determination of the pixel co-ordinates of the left and right wingtips in each of the *Wex* frames, and determination of the pixel co-ordinates of the head in each frame of the video sequence. While we have carried out all of these operations manually, they are tedious and time-consuming. Automated tracking and digitisation of the wingtips and the head in the video sequence can be incorporated as an additional ‘front end’ to the system, which we are currently exploring.
v) The technique delivers true altitude measurements only at each full wing extension. Altitudes at the intermediate frames are obtained by linear (or spline-based) interpolation. These interpolated heights can be combined with the digitized image position of the head in each frame to obtain a continuous, frame-by-frame 3D trajectory of the bird’s head. It is important to note that the X and Y positions of the bird’s head are tracked and reconstructed by using new information from *every* frame. The height interpolation should produce reasonably accurate results, provided that the bird’s altitude varies smoothly between successive wing extensions. This is very likely to be the case in cruising flight, but may not apply during flight through densely cluttered environments which may entail abrupt changes of altitude as well as variations in the wing kinematics.

Potential future applications of the method presented in this paper include:

i. Tracking of birds in natural outdoor environments by using an upward-facing wide-angle camera, as discussed briefly above. The species of the bird would have to be known, in order to use an estimate of its wingspan. However, even if the wingspan is not known, roll angles can continue to be computed, and the 3D trajectories of the head and thorax can be reconstructed in units of wingspan. This is demonstrated in Section C of the Supplementary Information. Thus, even if the wingspan of the bird is not known, it is possible to obtain several scale-invariant properties of the bird’s trajectory such as its shape, tortuosity, slope of ascent/descent and roll angle, as well as the timing and features of salient temporal events such as the onset of accelerations or decelerations, or the frequency of oscillatory movements. The method can also be applied to reconstruction and analysis of the flight trajectories of multiple birds (for example, in a flock). Again, even if the wingspan is not known, the spatial and temporal properties of the flock can be characterised by specifying inter-bird separations in wingspan units.
ii. Reconstruction of 3D flight trajectories of airplanes. In such an application, the 3D coordinates of the airplane and its roll angle can be estimated in every frame without any need for interpolation, because the wingspan is constant (as in a gliding bird). Again, the model of the aircraft would need to be known or identified, in order to use an estimate of its wingspan; otherwise, the aircraft’s trajectory can be specified in wingspan units.

## Acknowledgements

This work was funded by the ARC Centre of Excellence in Vision Science (Grant CEO561903), ARC Discovery Grant DP 110103277, a Human Frontiers in Science Grant RGP0003/2013, a Queensland Premier’s Fellowship, and an ARC Distinguished Outstanding Researcher Award (DP140100914) to MVS.

## Author contributions

*MVS:* Conceptualization, data analysis, modeling, writing, and funding acquisition

*HD Vo:* Conduct of experiments, data collection and analysis, and writing

*I Schiffner:* Conduct of experiments, data collection and analysis, and writing

## Supplementary Information

### SECTION A

**Table S1.**
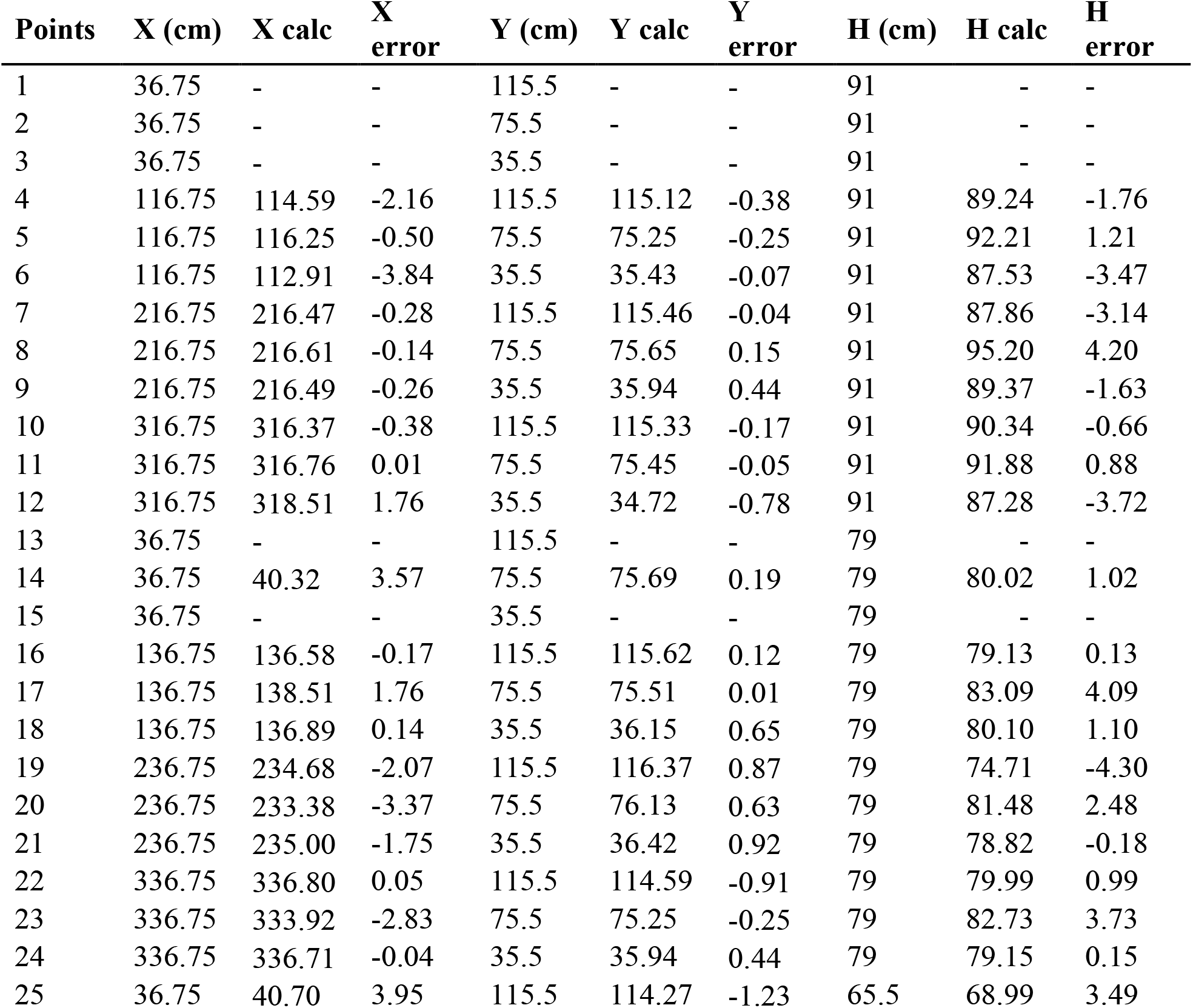
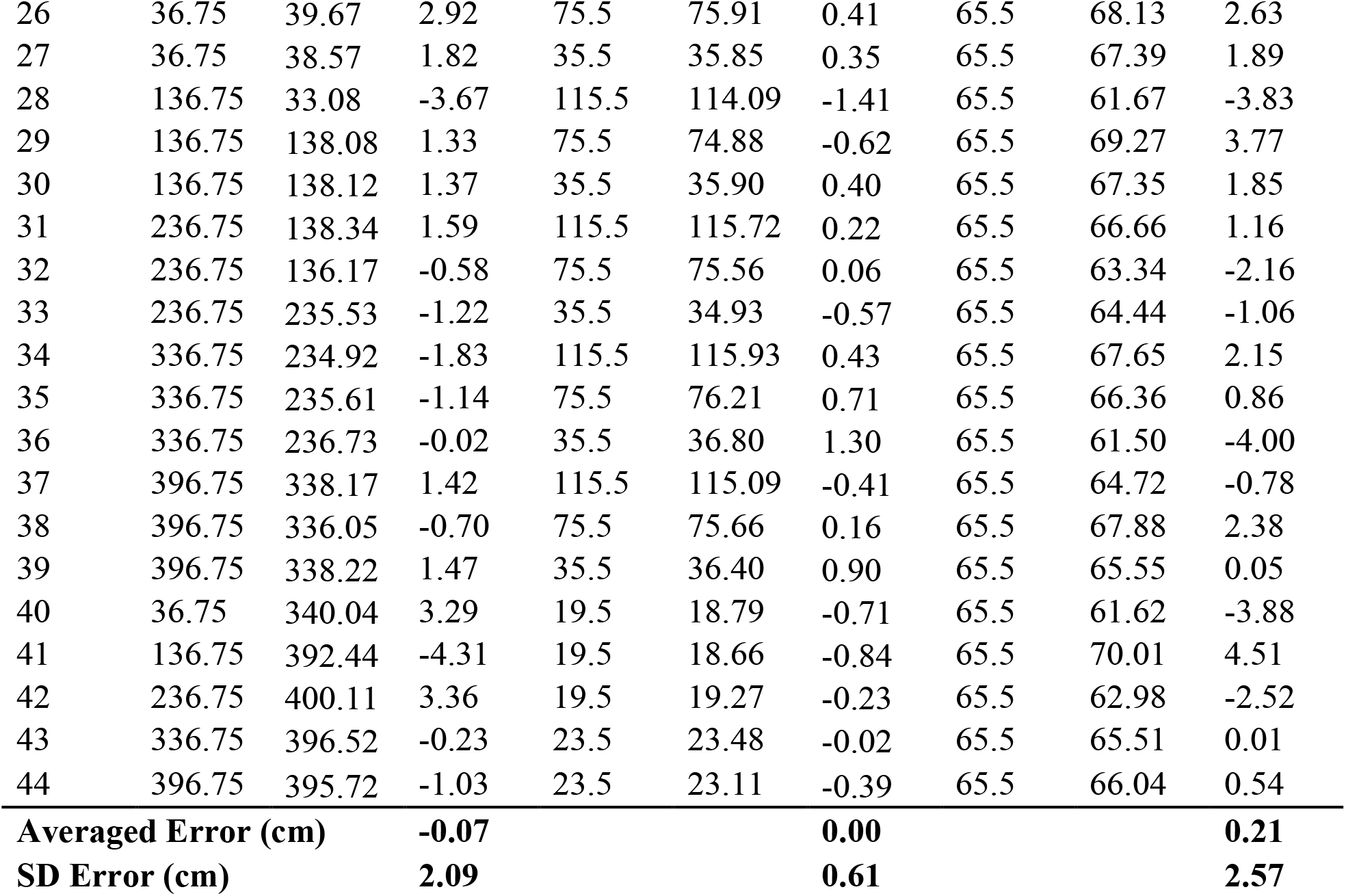
Test of accuracy of 3D head position measurements. X, Y and H represent (in cm) the true co-ordinates of a test target along the axial (length), width and height of the tunnel, respectively. X calc, Y calc and H calc are the calculated values of these co-ordinates, and X error, Y error and H error represent the respective errors. The standard deviation of the errors (SD) are given at the end of the table. The 5 missing measurements pertain to target positions whose floor projections fell outside the grid.

#### Results for bird Nemo

**Figure S1.**
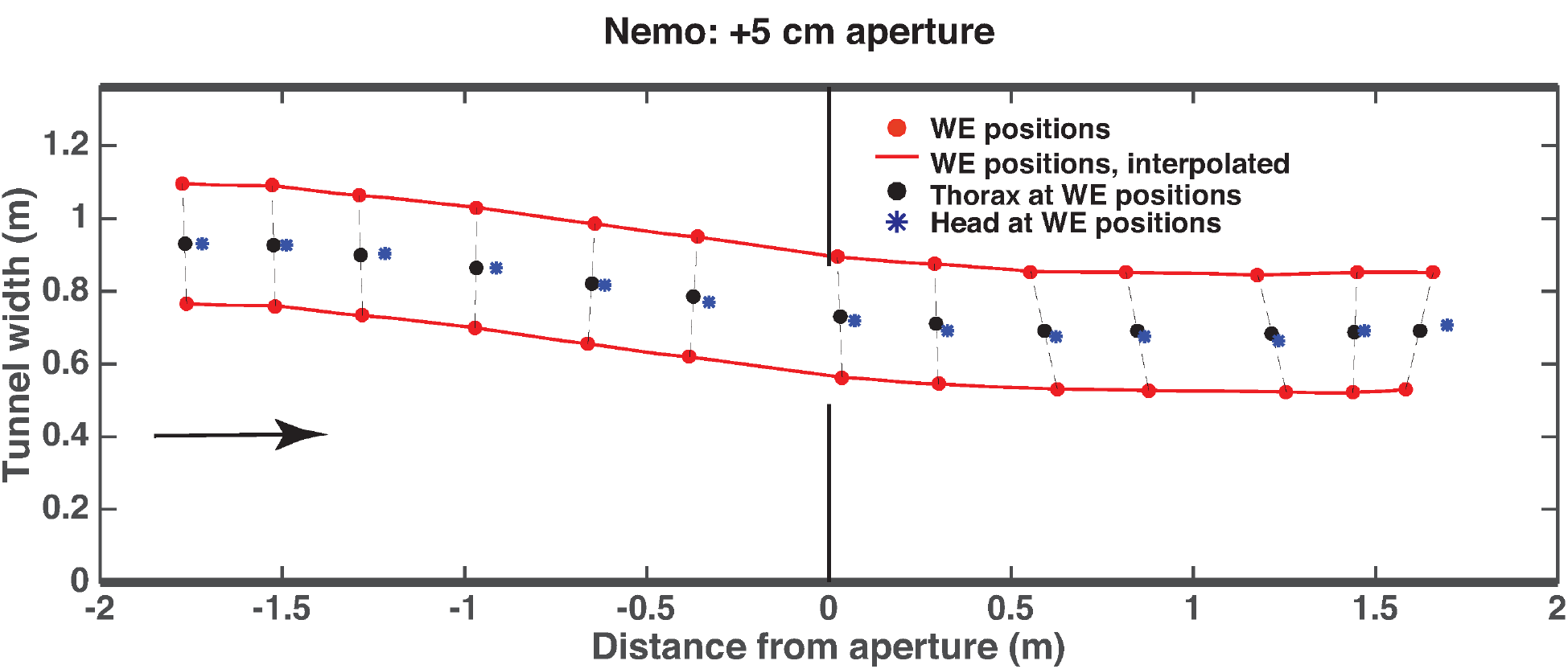
Plan view of a reconstructed flight of bird Nemo. In this example the wingspan of the bird (Nemo) is 33 cm and it flies through a 38 cm aperture, which is 5 cm wider than the wingspan. Details are as in Figure 5.

**Figure S2.**
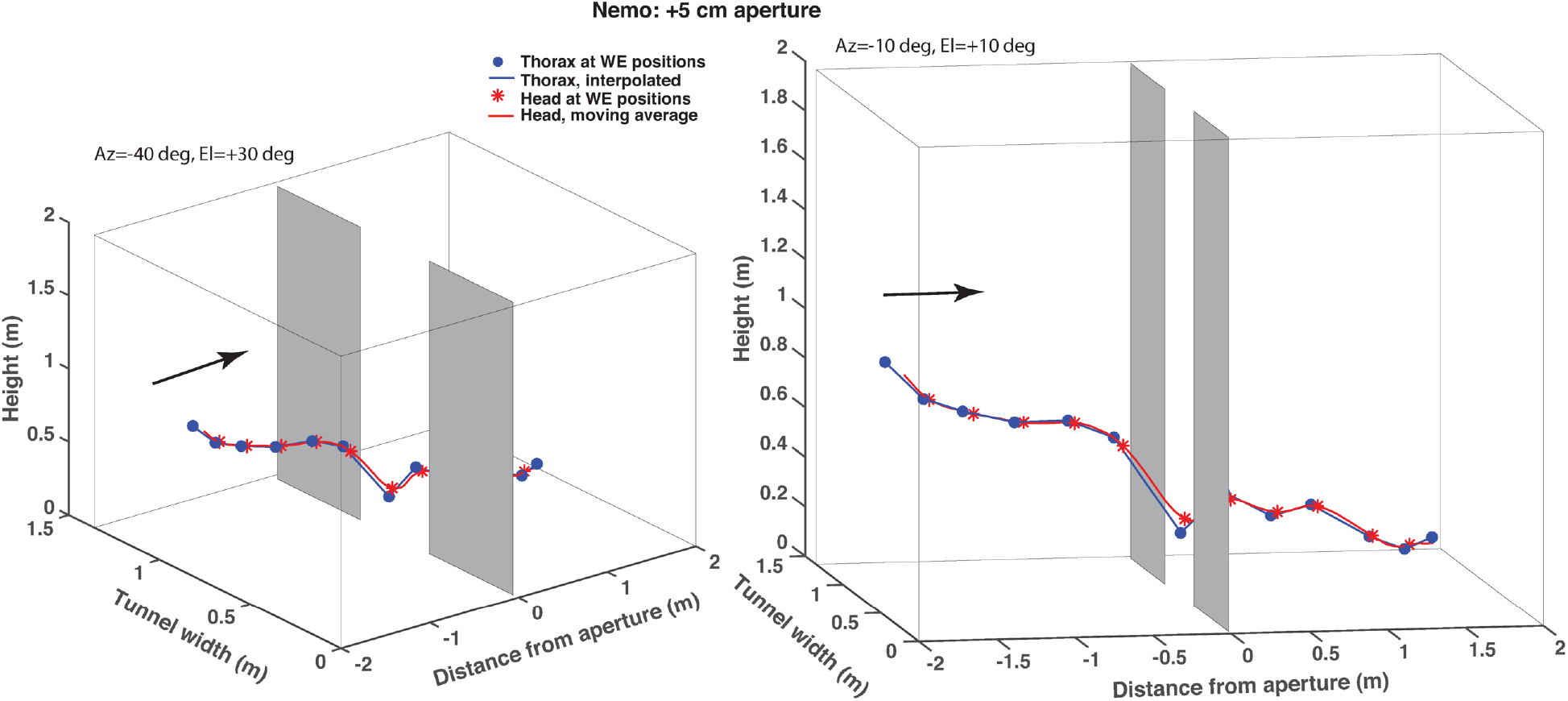
Two 3D views of the trajectory shown in Figure S1, in which Nemo flies through an aperture that is 5 cm wider than its wingspan. Details are as in Figure 6.

**Figure S3.**
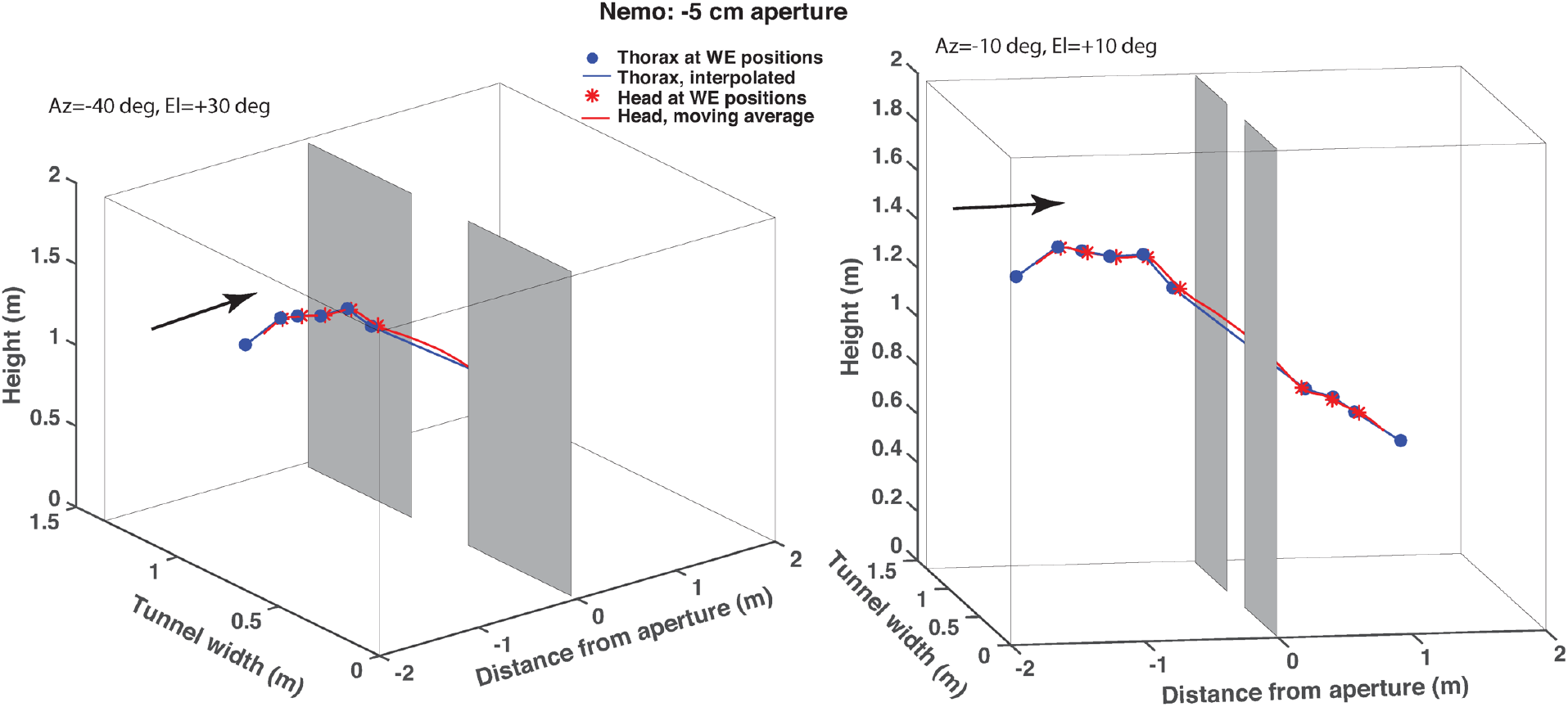
Two 3D views of a trajectory of bird Nemo during flight through an aperture that is 5 cm narrower than its wingspan. Details are as in Figure 6.

**Figure S4.**
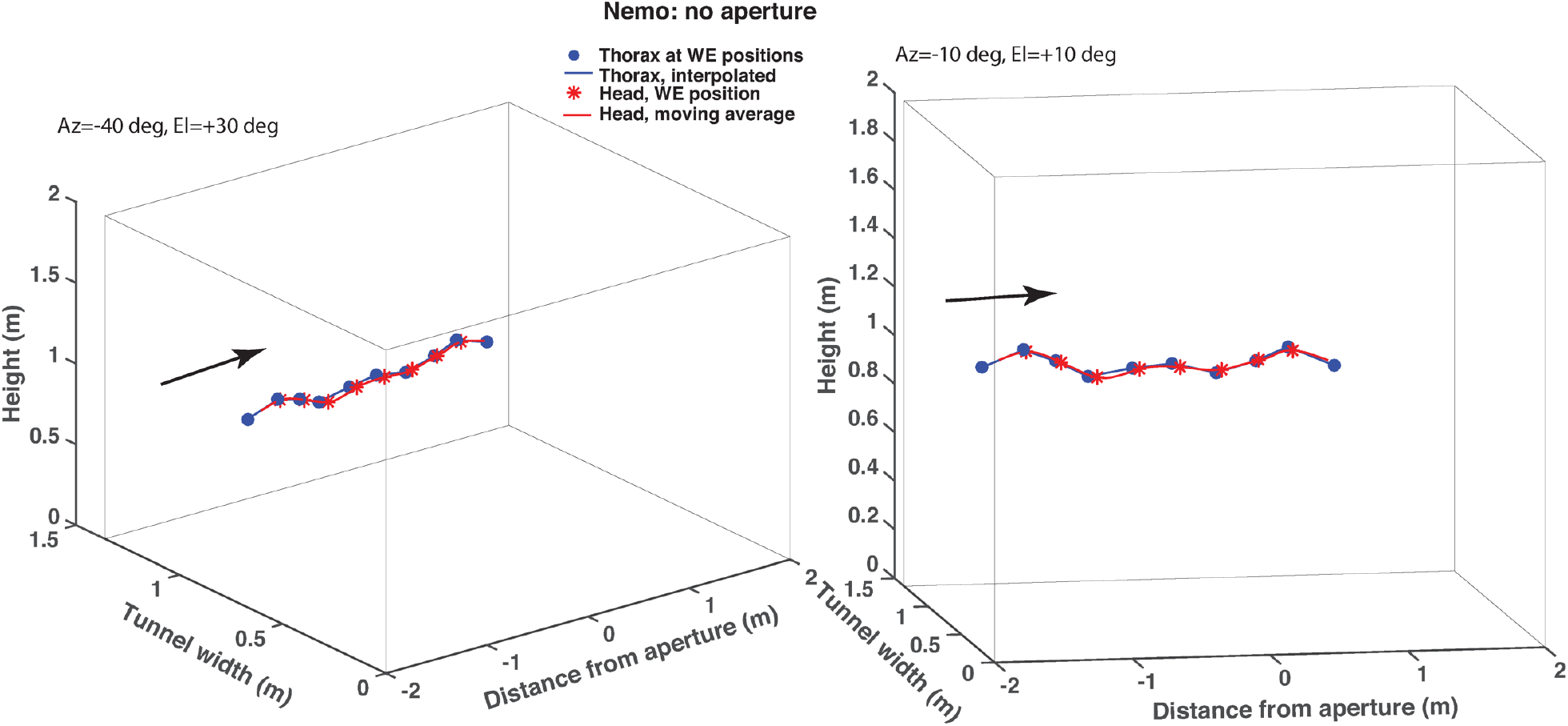
Two 3D views of a trajectory of bird Nemo during flight through a tunnel which carries no aperture. Details are as in Figure 6.

**Figure S5.**
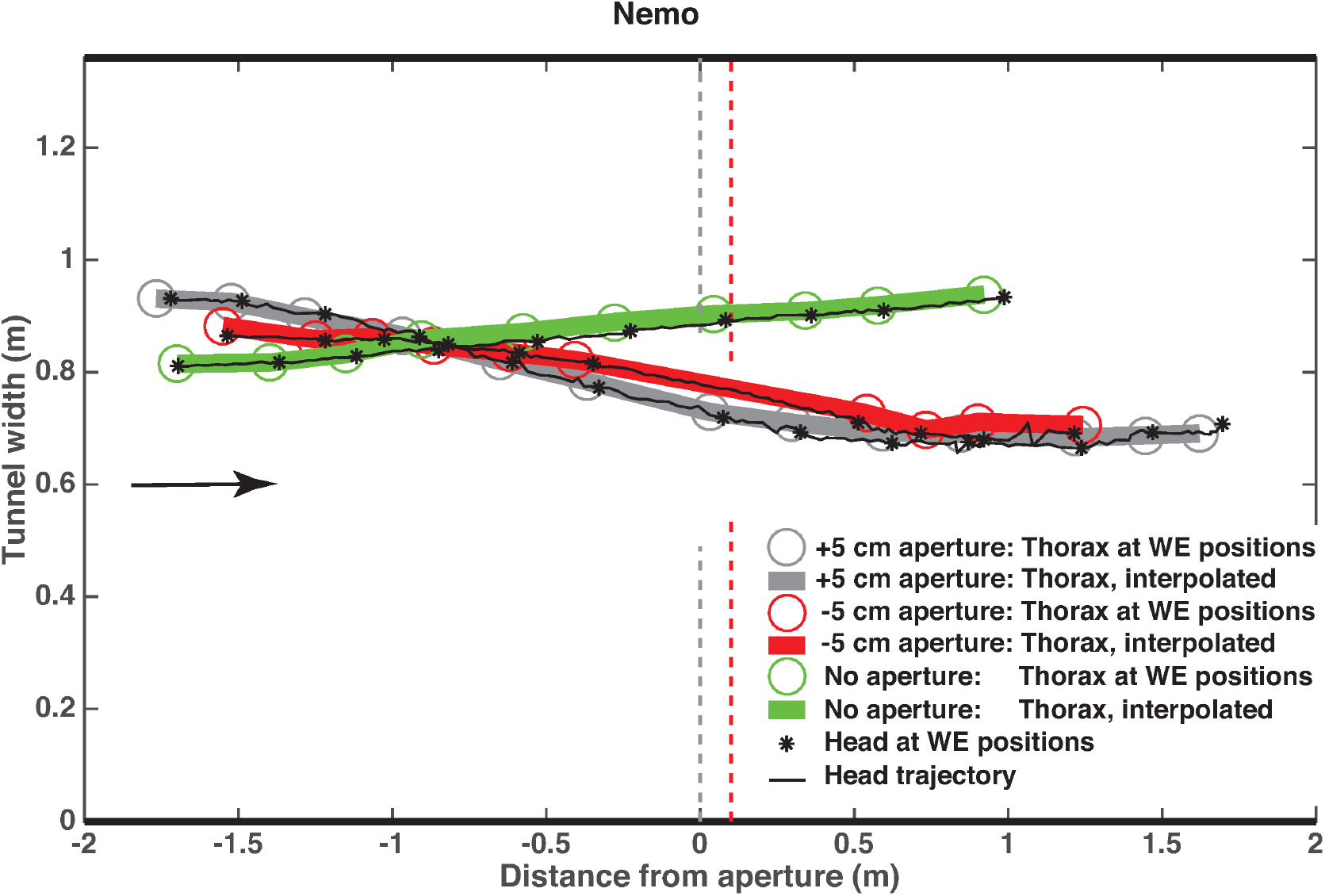
Plan views of the reconstructed 3D trajectories for the narrow aperture condition (red), the wide aperture condition (grey) and the no aperture condition (green), for bird Nemo. Details are as in Figure 9.

**Figure S6.**
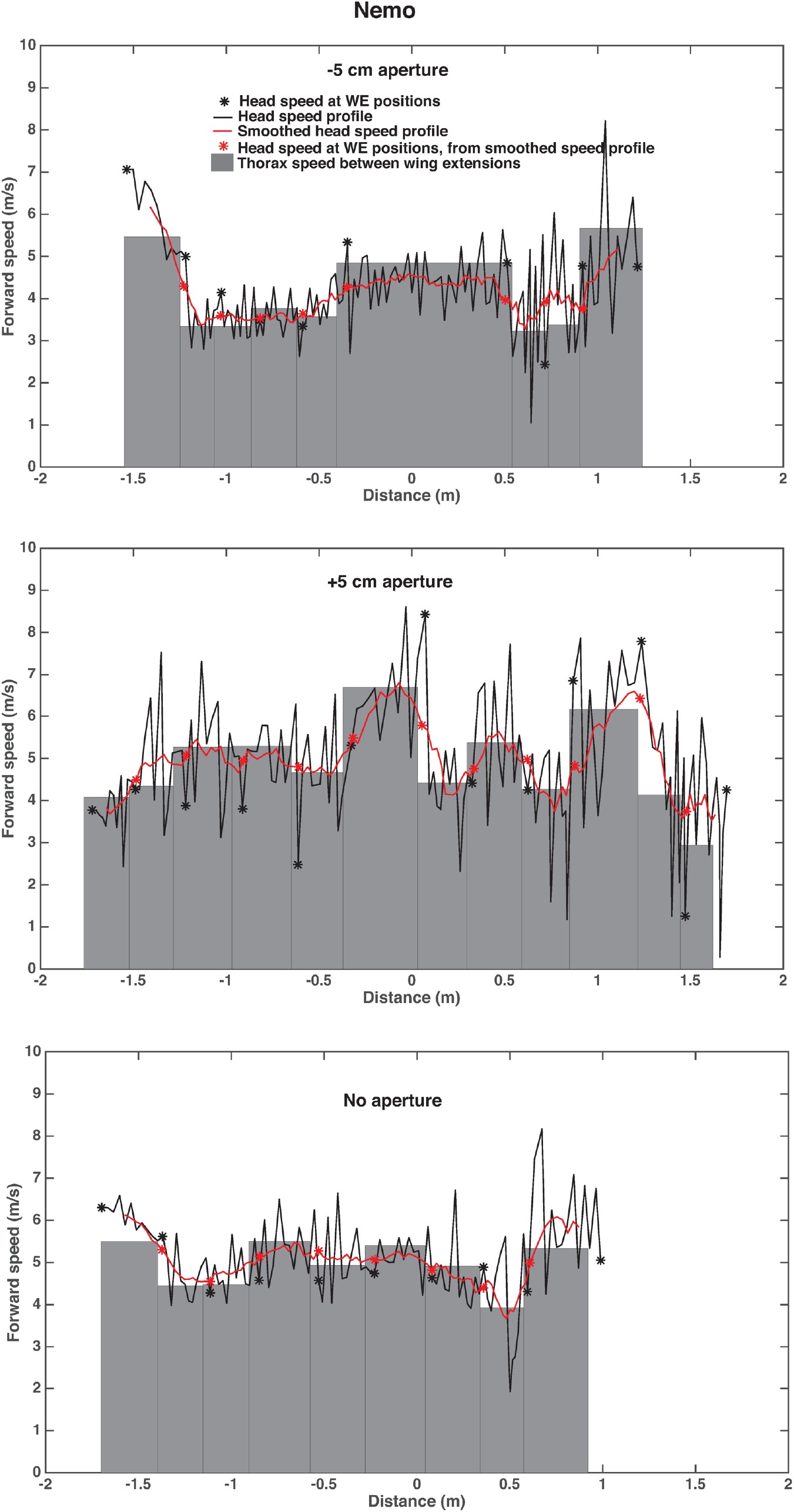
Forward speed profile of bird Nemo during flight through the narrow aperture (top panel), the wide aperture (middle panel), and the empty tunnel (bottom panel). Details are as in Figure 10.

### Supplementary Videos

Two examples of the flights of bird *Four,* as captured by the overhead video camera, are shown in videos SV1 and SV2. In video SV1 the bird passes through an aperture that is 5 cm wider than its wingspan (+5 cm) without closing its wings. In SV2, the bird passes through an aperture that is 5cm narrower than its wingspan (−5 cm), and closes its wings before entering the aperture. The calibration grid is not visible in these videos because its construction and camera calibration were carried out after the flights were filmed.

SV1: https://drive.google.com/file/d/1kKwyJ8IJtk3q7357YKiwodz-Ydlmya0M/view?usp=sharing

SV2: https://drive.google.com/file/d/1n3FjqKH_oMk5Wfsb_xqQQS7zVJ9BhCkd/view?usp=sharing

## SECTION B

### Derivation of extended calculation

Figure S7 illustrates the general case in which a bird, at an arbitrary 3D position in the arena, is filmed by the overhead camera. The floor caries a calibration grid (not shown). The bird is shown rolling to the left (the left wingtip is lower than the right). We wish to determine the height (h) of the bird above the floor, and the roll angle (a). R and L denote the positions of the right and left wingtips when the wings are fully extended; O denotes the thorax point, as defined in the main text. X, Y and Z denote the projections of the images of R, L and O on the floor. These are locations determined by grid interpolation, as described in Section 2.1 of the main text. U, the intersection of the camera axis on the floor, is the origin of the 3D coordinate system with coordinates (0,0,0). Since X,Y and Z are on the floor, their z-coordinates are zero. Other points and geometrical variables are defined in the figure. Figure S8 shows a perpendicular view of the plane PXY, including definitions of additional variables (including the angles ϕ_1_, ϕ_2_) that are relevant to the calculation.

**Figure S7.**
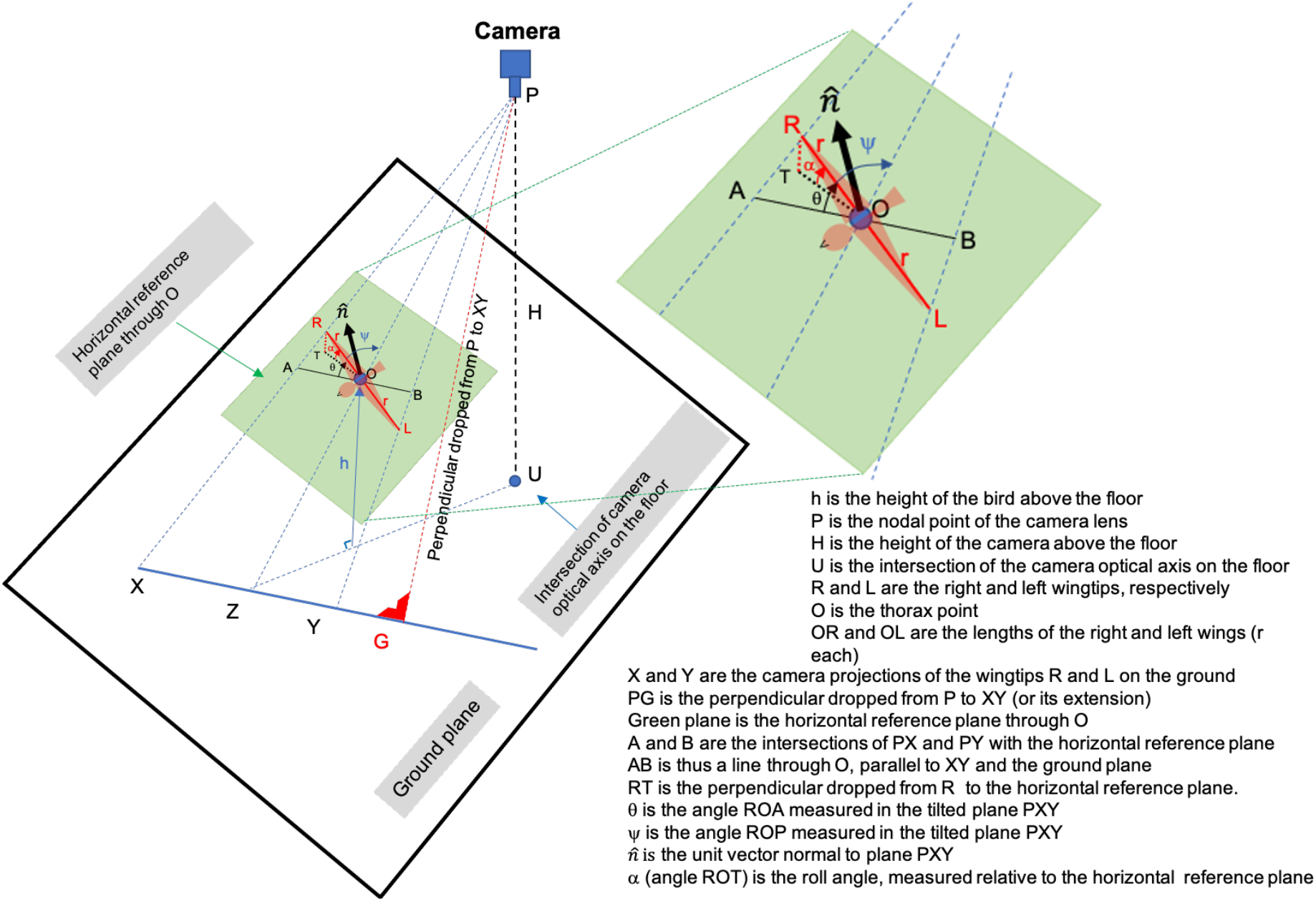
Geometry pertaining to calculation of the height and the roll angle of a bird at an arbitrary 3D location.

**Figure S8.**
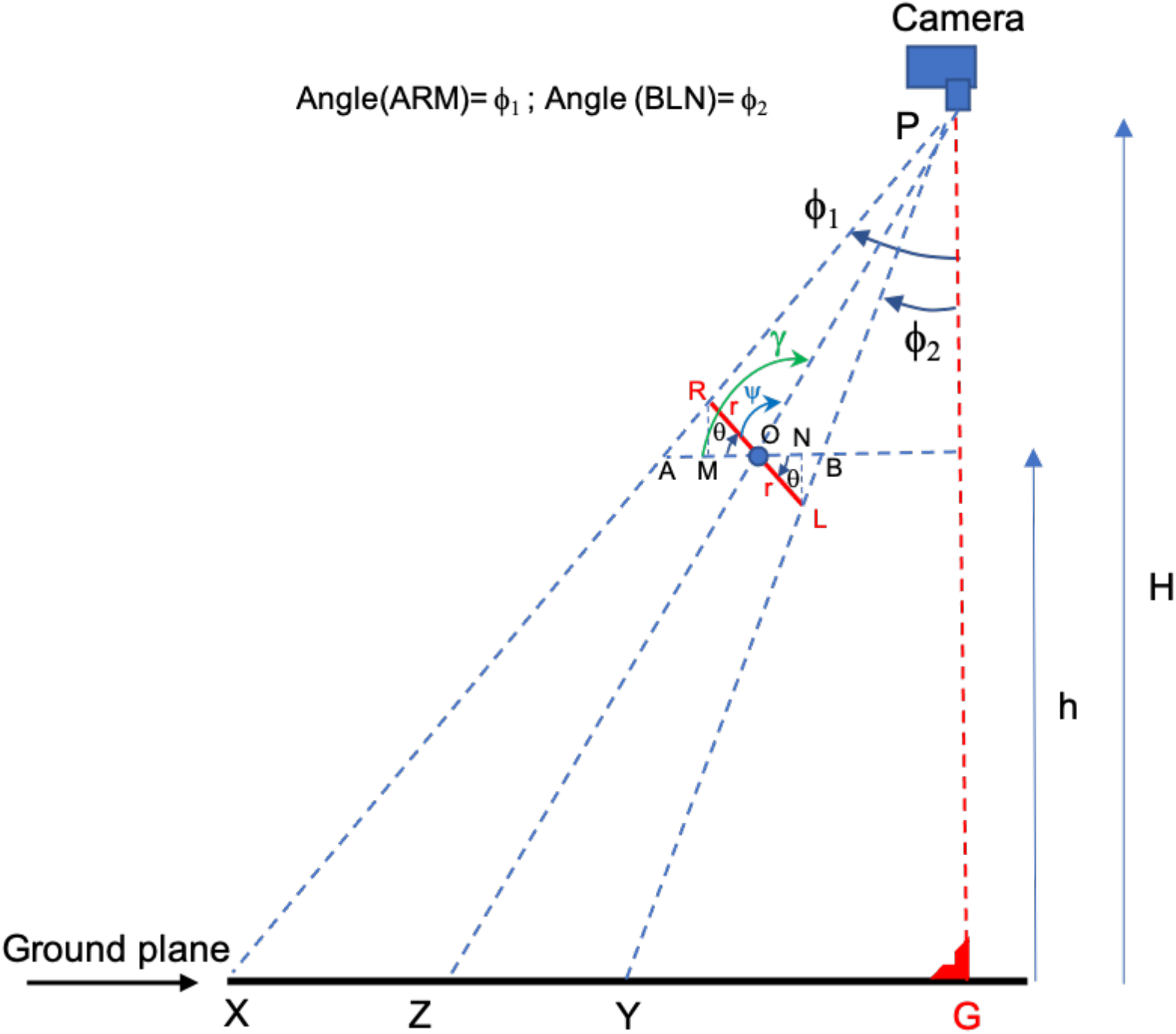
Perpendicular view of the plane PXY in Figure S7. Details in text.

The calculation is performed in three stages. First, we determine θ, which is the angle ROA measured in the plane PXY (See Figures S7, S8; Note that this angle is different from the roll angle a). Next, we use the calculated value of θ to determine the height (*h*) of the bird (this is the height above the floor of the thorax point O). Finally, we use the values of θ and *h* to calculate the roll angle a, which is the angle ROT in Fig S7.

#### Stage 1: Calculation of θ

Since X, Y and Z are the camera-projected points of the right and left wingtips (R,L) and the thorax point (O) on the floor, their floor coordinates, and hence the distances [XY], [XZ] and [YZ] can be determined.

Next, we determine the location of the point G, which is the intersection of the perpendicular dropped from P to the line connecting X and Y (or its extension). The location of G relative to X and Y can be determined by calculating the 3D vectors 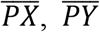 and 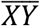, and the unit vector in the direction of 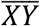, which we denote by 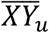. The length [GX] is the projection of 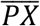 on 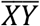, which is given by 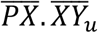 where (.) denotes the vector dot product. Similarly, the length [GY] is given by 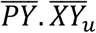.

From the geometry of Figure S8, it is clear that angle(ARM) = ϕ_1_, and angle(BLN) = ϕ_2_.

Therefore,

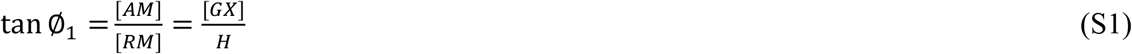

and

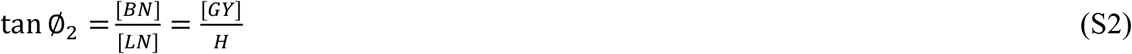

Since [GX], [GY] and H are known, the angles ϕ_1_ and ϕ_2_ can be evaluated from (S1) and (S2).

From Figure S8, we have

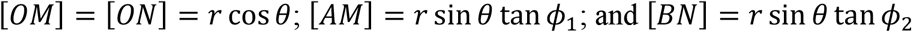

We can thus write

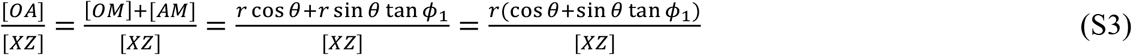

and

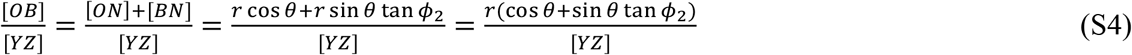

From triangle similarity, we have 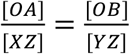

Equating (S3) and (S4), we obtain

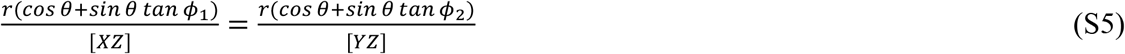

or

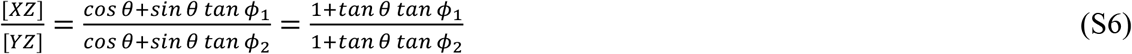

where [XZ] and [YZ] are known (see above), and ϕ_1_ and ϕ_2_ have been determined from (S1) and (S2) above.

Denoting the ratio 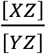 by Q (which is now known), we can solve for *tan θ* from (S6) to obtain

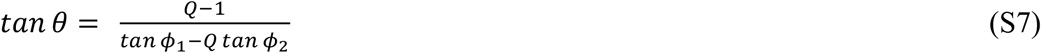

Since ϕ_1_ and ϕ_2_ are known from (S1) and (S2), (S7) can be used to calculate *tanθ,* and hence θ.

#### Stage 2: Calculation of bird height (h)

Referring to Figure S8, since triangles PAB and PXY are similar, we can write

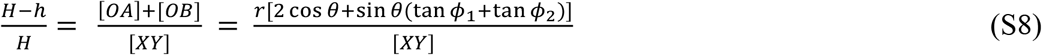

H and *r* are known, and ϕ_1_, ϕ_2_ and θ have been calculated from (S1), (S2) and (S7), respectively. Hence, we can solve for h from (S8) to obtain

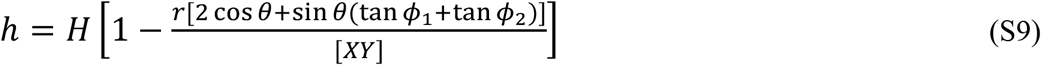

We note that ϕ_2_ will be positive or negative according whether Y is to the left or the right of G. (PG is the perpendicular dropped from P to the line XY, or its extension). If [XG] > [XY], X and Y are on the same side of G. If [XG] < [XY], X and Y are on opposite sides of G. In our example the negative value of ϕ_2_ is used when solving for θ in (S7), and for *h* in (S9).

Once the height of the thorax point (*h*) is known, the 3D coordinates of the thorax point and the head can be calculated using the grid interpolation procedure described in Section 2.1 of the main text.

#### Stage 3: Calculation of roll angle (α)

Referring to Figure S7 we note that U, the point on the floor directly beneath the camera, is the origin of the 3D co-ordinate system, with co-ordinates (0,0,0). P, the nodal point of the camera lens, has co-ordinates (0,0,H). We also know the 3D co-ordinates of X, Y and Z, which have been projected from the camera image to the floor. (As these points are on the floor, they will each have a *z* coordinate of zero). Using this information we calculate the 3D coordinates of A, which we denote by the 3D vector 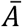, as follows:

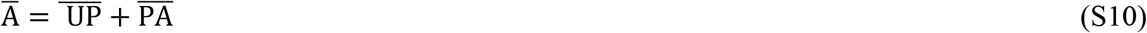

From triangle similarity, we have 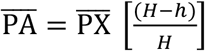. Hence (S10) can be expressed as

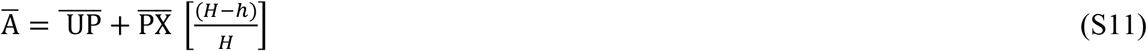

Similarly, the 3D coordinates of B and O are expressed by the vectors

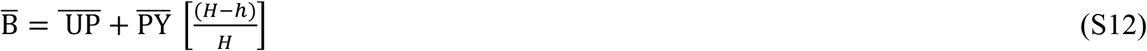

and

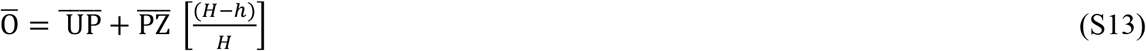

Next, we compute the 3D vectors 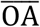 and 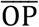 as

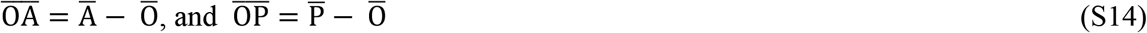

We can then use the unit vectors 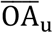 and 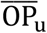, representing the directions of vectors 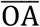 and 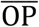, to compute the angle AOP, which we denote by *γ* (Figure S7):

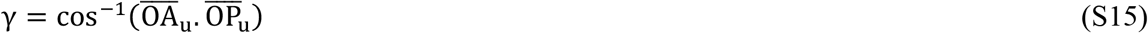

where (.) denotes the dot product.

From the angles AOR (*θ*) and AOP (g), calculated from (7) and (15), we can calculate the angle ROP, denoted by ψ in Figures S7 and S8, as

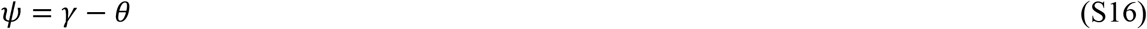

We can also use the unit vectors 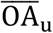 and 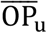 to determine the unit vector 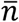, defining the normal to the plane POA (see Figure S7), as

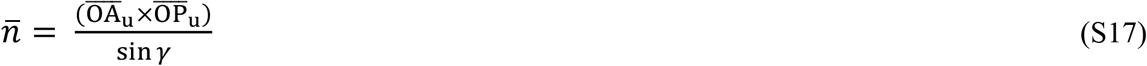

where (×) denotes the cross product.

To determine the roll angle of the bird, we need to determine the direction of the 3D vector 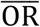 (Figure S7), which is the vector representing the direction of the right wingtip. We denote the unit vector in this direction by 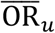. This unit vector can be determined from its relationship to the orientation of three other known vectors, as follows. First, since angle(AOR) = θ, we may write

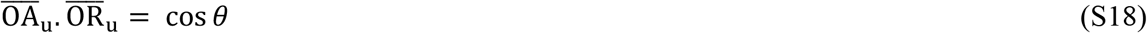

Second, since angle(ROP) = ψ, we may write

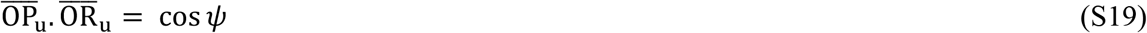

Finally, since 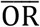 lies in the plane AOP, which is normal to 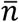, we may write

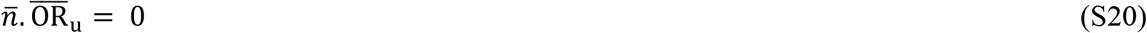

The 3D unit vectors 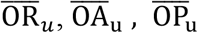 and 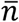 can be represented as

[OR_ux_ OR_uy_ OR_uz_ [OA_ux_ OA_uy_ OA_uz_], [OP_ux_ OP_uy_ OP_uz_], and [n_x_ n_y_ n_z_] where the suffixes *x, y* and *z* denote the *x, y* and *z* components of these vectors.

Equations (S18-S20) can thus be re-expressed as

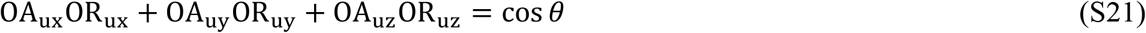

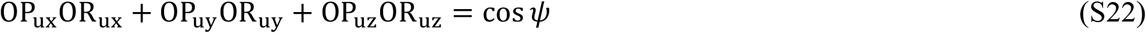

and

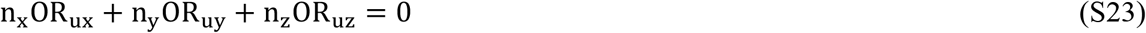

Equations (S21-S23) can be written in matrix form as

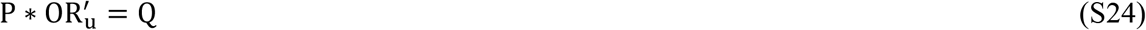

where (*) denotes matrix multiplication
and where

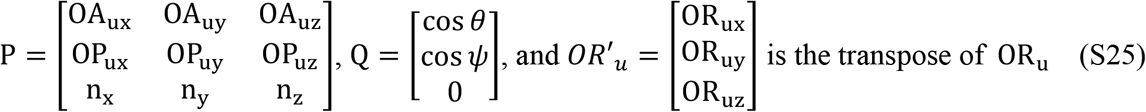

Since the elements of P and Q are known, 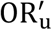 (and hence OR_u_) can be computed from (S24) using the matrix inverse of P:

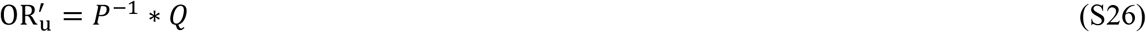

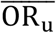 is the unit vector that defines the direction of 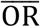, which is the 3D direction of the right wing when it is fully extended. The roll angle a is the angle between 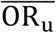 and the horizontal reference plane (see Figure S7). To compute this angle, we first calculate the angle β between 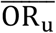 and the vertical (*z*) direction (not shown in Figure S7). Denoting the vertical direction by the unit vector 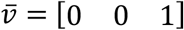, β can be calculated from

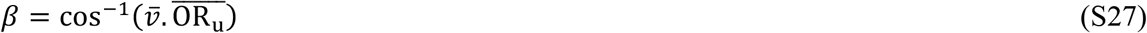

The roll angle a is then given by

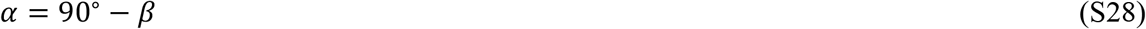

In our formulation the roll angle α is positive or negative according to whether the right wingtip is higher or lower than the left wingtip.

## SECTION C

### Reconstruction of 3D trajectories when the wingspan is unknown

Here we demonstrate that, even when the wingspan of the bird is not known, the 3D flight trajectory can be reconstructed in units of wingspan, regardless of whether or not the bird is rolling.

#### Zero-roll condition

Referring to the main text, equations (2), (3) and (6) can be used to express (8) as:

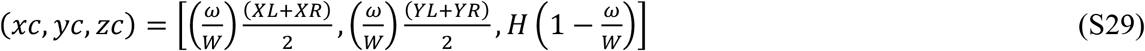

Denoting the depth of the bird below the ceiling by *h*’, we have 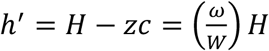.

Expressing the 3D co-ordinates of the center of the bird in terms of (*xc, yc, h’*), we can re-express (S29) as

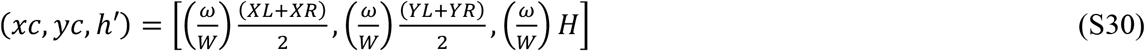

Thus, the 3D position of the bird (and its trajectory and velocity) can be specified in terms of the wingspan unit (ω), which acts as a scale factor.

#### Non-zero roll angle

If the roll angle is not zero, the angles θ and α can continue to be evaluated even when the wingspan is unknown, as is evident from equations (S7) and (S10 - S27). The depth (*h’*) of the thorax of the rolling bird, expressed in wingspan units (ω), can then be computed from equation (S9) as

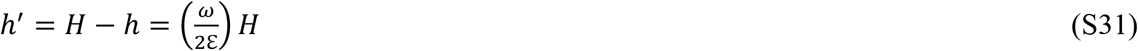

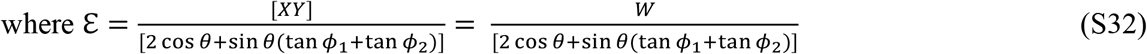

(When the roll angle θ is zero,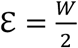 and (S31) reduces to 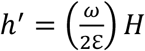, which coincides with the result shown in (S30)).

Once the height *h* of the thorax above the floor [*h* = (*H* – *h*′)] is known, the 3D coordinates of the thorax and the head can be evaluated (in wingspan units) using the grid interpolation procedure described in Section 2.1 of the main text:

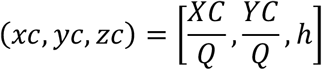

and

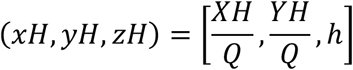

Thus, even if the wingspan of the bird is not known, it is possible to obtain several scale-invariant properties of the bird’s trajectory such as its shape, tortuosity, slope of ascent/descent and roll angle, as well as the timing and features of salient temporal events such as the onset of accelerations or decelerations, or the frequency of oscillatory movements.

